# Mechanism of duration perception in artificial brains suggests new model of attentional entrainment

**DOI:** 10.1101/870535

**Authors:** Ali Tehrani-Saleh, J. Devin McAuley, Christoph Adami

## Abstract

While cognitive theory has advanced several candidate frameworks to explain attentional entrainment, the neural basis for the temporal allocation of attention is unknown. Here we present a new model of attentional entrainment that is guided by empirical evidence obtained using a cohort of 50 artificial brains. These brains were evolved *in silico* to perform a duration judgement task similar to one where human subjects perform duration judgements in auditory oddball paradigms^1^. We found that the artificial brains display psychometric characteristics remarkably similar to those of human listeners, and also exhibit similar patterns of distortions of perception when presented with out-of-rhythm oddballs. A detailed analysis of mechanisms behind the duration distortion in the artificial brains suggests that their attention peaks at the end of the tone, which is inconsistent with previous attentional entrainment models. Instead, our extended model of entrainment emphasises increased attention to those aspects of the stimulus that the brain expects to be highly informative.

---

Our ability to deduce causation, to predict, infer, and forecast, are all linked to our perception of time. This activity of the brain refers to an inductive process that integrates information about the past and present to calculate the most likely future event (1). Without a doubt, this ability is key to an organism equipped with such a brain to survive and prosper, by predicting and deciphering events in the world (2, 3)–as well as the actions of other such organisms. A typical experimental procedure in the study of time perception is comparative duration judgement, in which subjects are asked to compare and judge the duration of events. Generally, duration judgements display the *scalar property*, which implies that the probability distribution of judgements is scale invariant (4). However, we do not perceive time objectively. Rather, the experience of temporal signals is highly subjective, and is influenced by non-temporal perception, attention, as well as memory (5, 6). An example of non-temporal perception is the saliency of a stimulus (how it stands out over a background), which may affect how it is perceived.

Attention is another variable that can shape time perception (7–11). Because our cognitive bandwidth is limited, we cannot pay attention to all sources of information equally (12). Rather, a sophisticated mechanism selects which stimuli are attended to, and how much attention is allocated to them. A central hypothesis is that the more attention is devoted to the duration of an event, the longer it is perceived to last (7–11). Proposed models of time perception such as Scalar Expectancy Theory (SET) (13) that support this hypothesis usually assume that duration perception is performed with some sort of internal clock (4, 13, 14). In that model, the onset of an event triggers a switch that starts measuring the accumulation of pulses generated by a *pacemaker*, and triggers the stop switch at the end of the event. The effective rate of pulse accumulation, in turn, is modulated by the attention given to the stimulus.

In SET, the amount of attention allocated to the stimulus is uniformly distributed in time. By contrast, in models such as Dynamic Attending Theory (DAT) (15–17) the temporal structure of the signal within which the stimulus is embedded may increase or decrease levels of attention in time. In particular, rhythmic backgrounds can *entrain* the brain so that it expects stimuli to occur periodically, and leads to peaks and troughs of attention. Consequently, models of attentional entrainment based on DAT posit that attentional rhythms that are internal to the cognitive architecture are *synchronised* by external rhythms, so that the external stimuli can then lead to an enhanced processing of events that occur precisely when they are expected to occur (18–20). Previous studies have provided support for DAT and related entrainment models, for example by showing that events that occur at rhythmically expected time points can be discriminated more easily than those that occur unexpectedly (17, 19, 21–23). In a recent study, McAuley and Fromboluti provided additional support for DAT and related entrainment models by studying the role of attentional entrainment on event duration perception (24). In that work, they used an auditory oddball paradigm in which a deviant tone (oddball) is embedded within a sequence of otherwise identical rhythmic tones (standard tones). Their results demonstrated that manipulations of oddball tone onset can lead to distortions in oddball tone duration perception. In particular, they observed a systematic *underestimation* of the duration of oddball tones that came early with respect to the rhythm of the sequence, and an *overestimation* of oddball duration in trials where oddballs arrived late with respect the rhythm of the sequence.

Interval timing models such as DAT and SET and their computational counterparts usually take a top-down approach by engineering networks of high-level computational components that describe behavioural/psychophysical data in duration perception (4, 15, 16, 25, 26) (see also references in (27–29)). Some studies have employed more elaborate models that consist of neuron-scale components (30, 31). Here, we take a bottom-up approach where evolution leads to a population of diverse computational networks (artificial brains) consisting of lower-level components. These brains may differ in their components and possibly in their behaviours (higher level computations). These modern computational methods have opened a new path towards understanding perception: the recreation, *in silico*, of neural circuitry that implements behaviour similar to human performance. While this capacity is still in its infancy and therefore can only emulate humans on fairly simple tasks (such as attentional entrainment), the usefulness of this tool for a future “experimental robotic psychology” (32, 33) is evident.

In this study, we use Darwinian evolution to create artificial digital brains, (also known as Markov Brains (34), see Methods), that are able to perform duration judgements in auditory oddball paradigms1. Markov Brains are networks of variables with discrete states that undergo transitions evoked by sensory, behavioural, or internal states, and capable of stochastic decisions. As such, they are abstract representations of micro-circuit cortical models (35), except that their dynamics is not programmed.

We run 50 replicates of the evolutionary experiment (i.e., 50 different populations) and from each pick the best-performing Brain. These evolved Brains display behavioural characteristics that are similar to human subjects: for example, their discrimination thresholds satisfy Weber’s law. In fact, these 50 Brains can be thought of as participants in a cognitive experiment. We then test these Brains against auditory oddball paradigms that they have never experienced before, in which the oddball tone may come early or late with respect to the rhythm of the sequence (similar to the first series of experiments in Ref. (24)). The evolved Brains show distortions in perception of early/late tones similar to what was reported in human subjects (24). We then analyse the algorithms and computations involved in duration judgement in order to discover how these algorithms result in systematic distortions in perception of early/late oddballs.

Our findings demonstrate that the computations involved in duration judgements and distortions is quite different from existing time perception theories such as scalar expectancy theory (SET) or dynamic attending theory (DAT), and suggest a new theory of perception in which attention to uncertain parts of the stimuli plays the central role, whereas predictable parts require less attention (i.e., less processing) because they are expected (36). This is consistent with recent findings that predictability of stimuli results in more rapid recognition (37). We close with speculations that suggest a broader view in which all cognitive processing can be understood in terms of context-dependent prediction algorithms that pay attention only to those parts of the signal that are predicted to have the highest uncertainty, and are therefore likely to be informative.

## Results

We evolve Markov Brains that are capable of duration judgements of an oddball tone placed in a rhythmic sequence of identical tones (standard tones) with a variety of standard tone durations and inter-onset-intervals (IOI) (Fig. 1 shows a schematic of the auditory oddball paradigm). We ran 50 replicates of the evolution experiment for 2,000 generations and from each population picked the Brain with the highest performance at the end of each run. The best performing Brains in all 50 populations gain 98.0% fitness on average (see Supplementary Fig. 1).

**Fig. 1.**
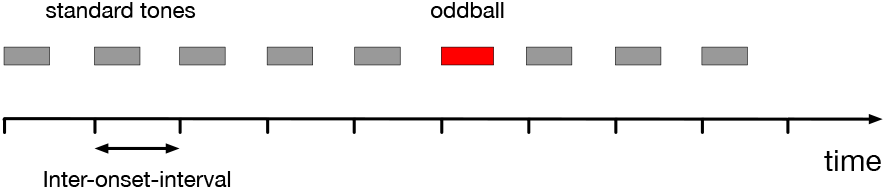
A schematic of the auditory oddball paradigm in which an oddball tone is placed within a rhythmic sequence of tones, i.e., standard tones. Standard tones are shown as grey blocks and the oddball tone is shown as a red block. Oddball tone duration may be longer or shorter than the standard tones.

### Discrimination thresholds of evolved Markov Brains comply with Weber’s law

We used average responses of the evolved Brains to generate psychometric curves as follows. For each (IOI, standard tone) we averaged the decision responses of 50 evolved Brains. Using these averaged responses, we generated psychometric curves corresponding to each standard tone as prescribed by (38) and calculated the point of subjective equality (PSE) and just noticeable difference (JND). The PSE measures the duration for which Markov Brains respond longer (or shorter) 50% of the time which, in essence, marks the duration of the oddball that is perceived to be equal to the standard tone. The JND measures the sensitivity of the discrimination, or discrimination threshold, for a standard tone. In other words, the JND represents the slope of the psychometric curve, where steeper slopes show higher discrimination sensitivity or lower discrimination threshold. The PSE reflects the accuracy of the perception while the JND indicates its precision. The values of PSE, JND, and their standard deviations are presented for all inter-onset-intervals and standard tones in Table 1.

**Table 1.**
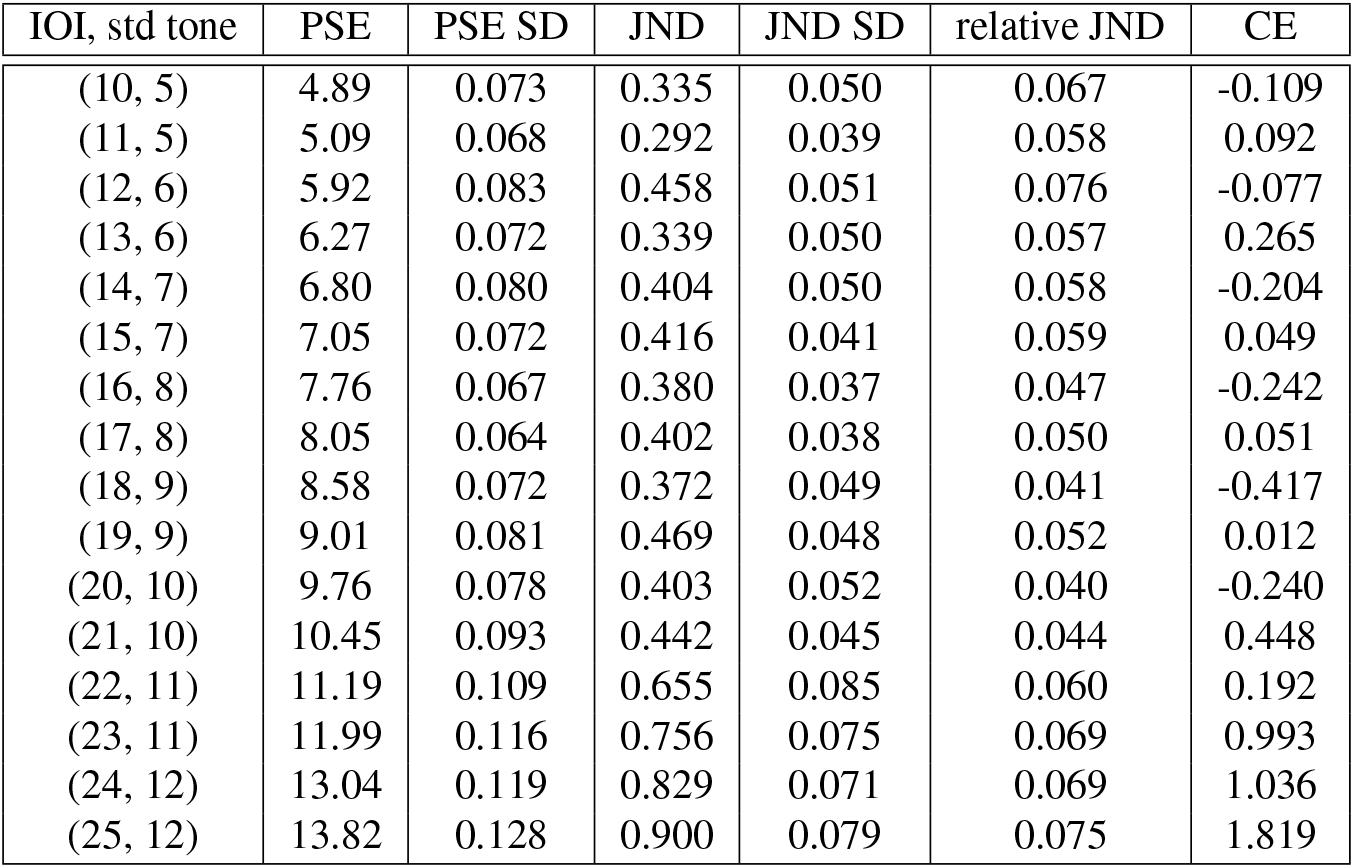
This table contains point of subjective equality (PSE), just noticeable difference (JND), and their standard deviations (SD), as well as relative JNDs, and constant error (CE) of on-time oddballs for all inter-onset-intervals, standard tones. Responses are averaged across all 50 Brains to generate psychometric curves.

| IOI, std tone | PSE | PSE SD | JND | JND SD | relative JND | CE |
| --- | --- | --- | --- | --- | --- | --- |
| (10, 5) | 4.89 | 0.073 | 0.335 | 0.050 | 0.067 | -0.109 |
| (11, 5) | 5.09 | 0.068 | 0.292 | 0.039 | 0.058 | 0.092 |
| (12, 6) | 5.92 | 0.083 | 0.458 | 0.051 | 0.076 | -0.077 |
| (13, 6) | 6.27 | 0.072 | 0.339 | 0.050 | 0.057 | 0.265 |
| (14, 7) | 6.80 | 0.080 | 0.404 | 0.050 | 0.058 | -0.204 |
| (15, 7) | 7.05 | 0.072 | 0.416 | 0.041 | 0.059 | 0.049 |
| (16, 8) | 7.76 | 0.067 | 0.380 | 0.037 | 0.047 | -0.242 |
| (17, 8) | 8.05 | 0.064 | 0.402 | 0.038 | 0.050 | 0.051 |
| (18, 9) | 8.58 | 0.072 | 0.372 | 0.049 | 0.041 | -0.417 |
| (19, 9) | 9.01 | 0.081 | 0.469 | 0.048 | 0.052 | 0.012 |
| (20, 10) | 9.76 | 0.078 | 0.403 | 0.052 | 0.040 | -0.240 |
| (21, 10) | 10.45 | 0.093 | 0.442 | 0.045 | 0.044 | 0.448 |
| (22, 11) | 11.19 | 0.109 | 0.655 | 0.085 | 0.060 | 0.192 |
| (23, 11) | 11.99 | 0.116 | 0.756 | 0.075 | 0.069 | 0.993 |
| (24, 12) | 13.04 | 0.119 | 0.829 | 0.071 | 0.069 | 1.036 |
| (25, 12) | 13.82 | 0.128 | 0.900 | 0.079 | 0.075 | 1.819 |

According to Weber’s law (39), the discrimination threshold (e.g., the JND) varies in proportion to the standard stimulus; therefore, the values of relative JND, defined as 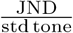, should remain constant. Getty showed that empirical results of duration perception in the range of 80 msec to 2 seconds is explained very well with Weber’s law (40). Fig. 2A shows the psychometric curves generated from the averaged responses of all 50 Brains for every (IOI, standard tone). In this figure, durations are normalised by standard tone. Psychometric curves for different standard tones overlap, which shows that relative JNDs in all these trials are similar and confirms that they are in accordance with Weber’s law. Fig. 2B shows relative JNDs as a function of standard tones. All relative JNDs are in the range between 0.04 and 0.07 with mean=0.06 and standard deviation=0.01, similar to the values found in (40). The difference between PSE and the standard tone, also known as the constant error (CE), shows the deviation of perceived duration of tone from its actual duration. The values of CE are shown for every (IOI, standard tone) in Fig. 2C and we observe that for longer IOIs, CE values start to deviate slightly from zero. This deviation in PSE values for longer tone durations is also observed in human subjects (40). However, this deviation of CEs for longer tones in Markov Brains was different from human subjects in that CE values in human subjects start decreasing for longer durations (they are negative) whereas in Markov Brains CE values increase (they are positive). This difference can be attributed to the fact that in the experiments described in Ref. (40) subjects do not receive any feedback about their performance duration judgements whereas Darwinian evolution provides feedback implicitly via selection. The mechanisms behind the distortion in duration perception in longer IOIs are explained in more detail in Supplementary Information.

**Fig. 2.**
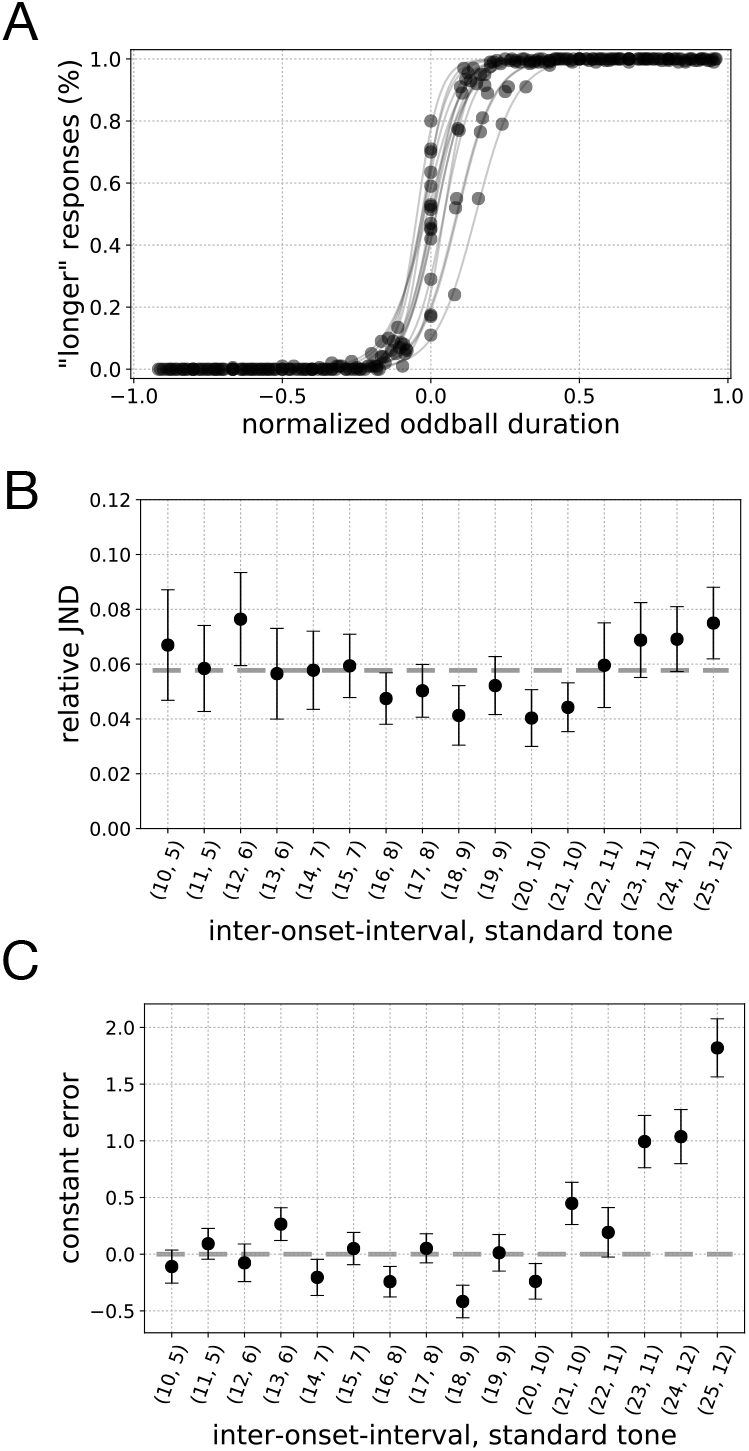
(A) Psychometric curves generated from averaged responses of 50 evolved Brains for every inter-onset-interval, standard tone. Oddball durations on the *x*-axis are normalised by standard tone to lie in the range (−1, 1). (B) Relative JND values and their 95% confidence interval as a function of inter-onset-interval, standard tone. Dashed line shows the average value of relative JNDs. (C) Constant errors, the difference between PSE and standard tone, and their 95% confidence interval as a function of inter-onset-interval, standard tone. Dashed line shows CE=0.

### Evolved Brains show systematic duration perception distortion patterns similar to human subjects

In the next step, we tested evolved Markov Brains with stimuli that they had never experienced during evolution, namely oddballs that arrive early or late with respect to the rhythm of the sequence of tones (termed “test trials”). In trials used during evolution (“training trials”), oddballs always occurred in sync with the rhythmic tone (on-time oddballs). These test trials included all possible oddball durations but also all possible oddball onsets, meaning oddballs were delayed or advanced as many time steps as possible as long as they did not interfere with the following or preceding tone. Then, we used the average response of 50 Brains to generate psychometric curves for early/late oddballs, and to calculate PSE values. We used PSE values to calculate the duration distortion factor (DDF), defined as the ratio of the point of objective equality (the standard tone) and the point of subjective equality (PSE).

Fig. 3 shows the DDF as a function of the onset of the oddball for all IOIs. In this plot, negative onset values stand for early oddballs and positive values of onset represent late oddballs. A DDF greater than one shows an overestimation of the duration of the oddball whereas a value less than unity reflects an underestimation of the duration of the oddball. Just as was observed with human subjects (24), the late oddballs are perceived as longer and the early oddballs are perceived as shorter compared to the standard tone. In addition, the more delayed (early) the oddball tone, the more its duration is overestimated (underestimated) compared to the standard tone, which is again consistent with results presented in experiment 2 of Ref. (24).

**Fig. 3.**
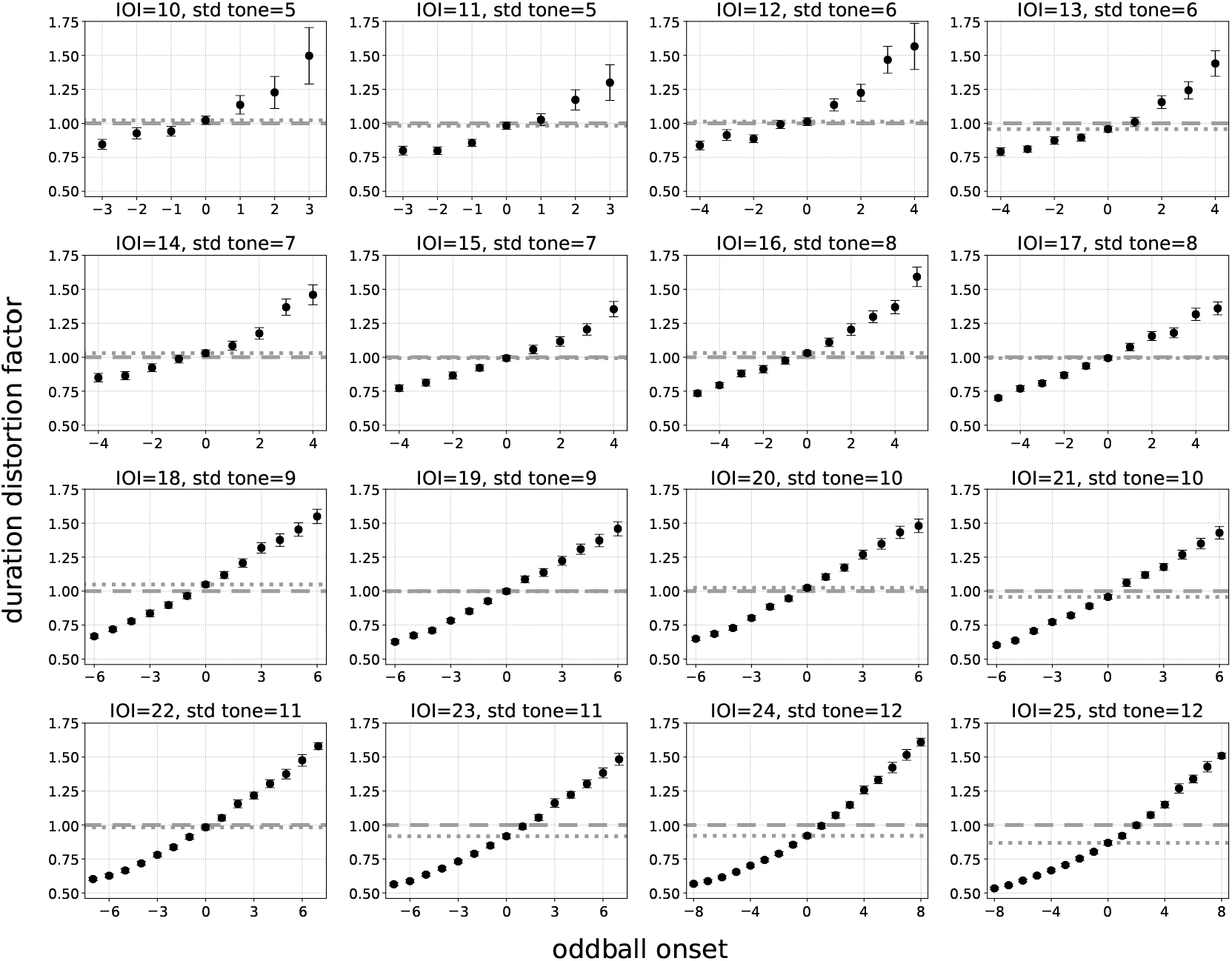
Duration distortion factors (DDF) and their 95% confidence interval as a function of the onset of the oddball for all IOI, standard tones. Negative onset values represent early oddballs and positive values of onset represent late oddballs. A DDF greater than 1 shows an overestimation of the duration of the oddball and DDF less than unity shows an underestimation of the duration of the oddball. The dashed line indicates DDF=1 and the dotted line shows DDF for on-time oddball tone.

### Algorithmic analysis of duration judgement task in Markov Brains

The logic circuits of evolved Markov Brains are complicated and defy analysis in terms of causal logic. As observed before, these networks turn out to be “epistemologically opaque” (41), in the sense that their evolved logic does not easily fit into the common logical narratives we are familiar with. Rather than focus on the Boolean logic of Markov Brains, we here focus on their state space (42, 43). In particular, we investigate the state transitions and how these transitions unfold in time, in order to discover the computations that are at the basis of the observed behaviour (44).

#### Temporal information about stimuli is encoded in sequences of Markov Brain states

Evolved Brains display periodic neural activation patterns in response to rhythmic auditory signals (this is, by definition, entrainment). These periodic neural firing patterns translate to loops in state transition diagrams (see Methods for more details on state transitions in Markov Brains). In each trial, the first few tones an evolved Brain listens to typically shift the Brain’s activation pattern towards a region in state space that is associated with this rhythm. More precisely, the opening tones transition the Brain to a sequence of states that form a loop in the state-to-state diagram, and the Brain remains in that loop as long as the stimulus is repeated. Fig. 4A shows an instance of a Markov Brain state transition diagram when listening to rhythmic tones with IOI=10 and standard tone=5 in the absence of an oddball. The state of the Brain is calculated from equation (2). Supplementary Movie 1 shows the state-to-state transitions as the Brain listens to a sequence of standard tones. This sequence of Brain states encodes the contextual information about the stimuli, that is, the sequence forms an internal representation of the rhythm and the standard tone. More importantly, this sequence produces an expectation of future inputs that enables the Brain to compare the input it has sensed with future inputs. In particular, when the Brain receives the oddball, it usually transitions out of this loop to follow a different trajectory in state space (see for example Fig. 4B) to judge the oddball duration, which is a compari-son mechanism between the standard tone (what is expected) and the oddball. Fig. 5 shows that in most of the trials (77.6% of the trials) Brains evolve loops of the same size as the period of the rhythmic tones (the IOI), but some Brains have loops that are multiples of the IOI. In this figure, the size of the each marker is proportional to the number of Brains that evolve a particular loop length in each IOI. Also, further analysis shows that in 93.6% of trials, evolved Brains transition out of these loops at the exact time point where there is a mismatch in oddball and standard tone.

**Fig. 4.**
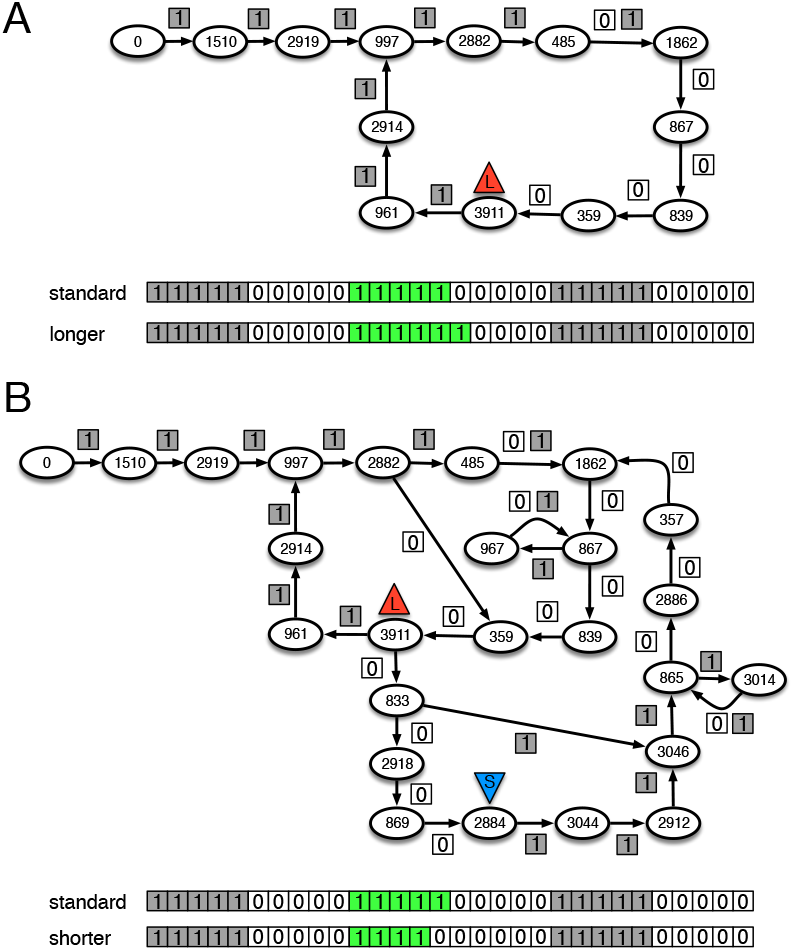
State-to-state transition diagram of a Markov Brain for IOI=10, and standard tone=5, with oddball tones of duration 5, 6 shown in (A) and 4 shown in (B). Before the stimulus starts, all neurons in the Brain are quiescent so the initial state of the Brain is 0. The stimulus presented to the Brain is a sequence of ones (representing the tone) followed by a sequence of zeros (denoting the intermediate silence). The stimulus at each time step is shown as the label of the transition arrow in the directed graph. The input sequence is shown for the standard and oddball sequences at the bottom of the state-to-state diagrams. (A) State-to-state transition diagram of a Markov Brain when exposed to a standard tone of length 5, as well as a longer oddball tone of length 6. This Brain judges an oddball tone of duration 6 by following the same sequence of states as the original loop, because the transition from state 485 to 1862 occurs irrespective of the sensory input value, 0 or 1. This Brain correctly issues the judgement “longer” from state 3911, indicated by the red triangle at the end of the time interval (see Supplementary Movie 1 and Supplementary Movie 2 for standard tone and longer oddball tone, respectively). (B) The state-to-state transition diagram of the same Brain when presented with a shorter oddball tone of length 4. The decision state is marked with a down-pointing blue triangle. Once the Brain is entrained to the rhythm of the stimulus, the shorter oddball throws the Brain out of this loop. The exit from the loop transitions this Brain into a different path. After four ones the Brain transitions to state 359 (instead of continuing to 485), and then continues along a path where it correctly judges the stimulus to be “shorter” in state 2884 (see also Supplementary Movie 3).

**Fig. 5.**
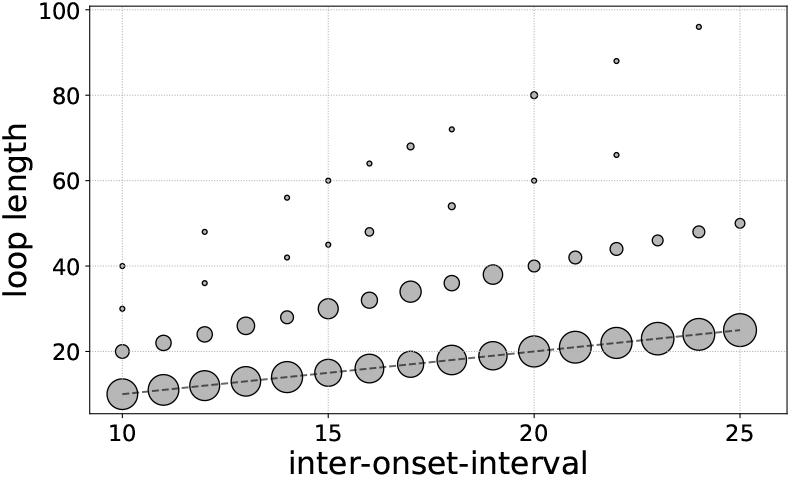
The distribution of loop sizes of 50 evolved Brain for each inter-onset-interval (IOI). The size of the markers is proportional to the number of Brains (out of 50) that evolve a particular loop length in each IOI. The dashed line shows the identity function.

### Algorithmic analysis of distortions in duration judgements: Experience and perception during misjudgements of early/late oddballs

The similarity of behavioural characteristics in the perception of event duration between Markov Brains and human subjects appears to imply a fundamental similarity between the underlying computations and algorithms. In the following, we present brief definitions of concepts such as attention, experience, and perception in terms of state transitions in deterministic finite-state machines that are later used in our analysis (in Methods we present more formal definitions of these concepts and the reasoning behind them).

1. Attention to a stimulus: When a Brain is in state *S_t_* and transitions to state *S*_*t*+1_ regardless of the stimulus (zero or one), we say the Brain *does not pay attention* to the input stimulus. More generally, a Brain pays less attention to an input stimulus or a sequence of stimuli if that input does not affect the state of the Brain later in the future state, *S_t+k_*.
2. Perception of a trial: The state of the Brain at the end of the oddball tone interval (when it issues the longer/shorter decision) is the Brain’s perception of the tone sequence.
3. Experience of the stimuli: The temporal sequence of Brain states when exposed to a sequence of input stimuli constitutes the Brain’s experience.

We first hypothesised that early or late oddball tones drive the Brain into states that they had never visited before (as these Brains had never previously experienced early or late oddball tones) and that these new states are responsible for misjudgements of early or late oddballs. When exposed to late or early oddballs, Brains visited on average 22.26 (SE=4.33) new states across 50 evolved Brains, approximately 32% of the number of states they visited during trials with on-time oddballs, which is 69.80 states on average (SE=5.07). We then tested how often these new states are decision states for the misjudgements of out-of-time oddball tones. Our tests show that in such misjudgements, the Brain state at decision time point is almost *never* a new state that has not appeared before (it happened in one test trial for one Brain out of 56,250 different test trials in all 50 Brains).

Given that during misjudgements of out-of-rhythm oddballs the decision state is a state that had previously occurred during evolution, we test whether there is any connection between Brain states during these misjudgements and Brain states in training trials. In other words, we investigate how the experience during a misjudgement relates to experiences the Brain had in its evolutionary history. In the next two sections, we address these questions by separately focusing on perception and experience of Markov Brains during misjudgements of out-of-rhythm oddball tones.

#### The onset of the tone does not alter a Brain’s perception of the tone

Our null hypothesis is that the perception of an out-of-rhythm oddball tone may be any one of the states that the Brain has traversed in training trials with equal probability. In any of these Brain states, the decision neuron will be either quiescent or firing, so we call the set of states with quiescent decision neuron “shorter-judging states” denoted as *S*_Sh_, and the set of states with firing decision neuron “longer-judging states” denoted as *S*_Lo_. Thus, the probability that a Brain at decision time is in any of the shorter-judging states, for example, is calculated by

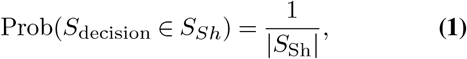

where |*S*_Sh_| is the cardinality of the set of shorter-judging states, and similarly, Prob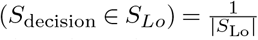.

We develop our alternative hypothesis that captures possible associations between experience and perception during misjudgement of out-of-rhythm oddballs and experiences and perceptions they had in training trials. In order to discover such possible associations, for any given misjudgement of early or late oddball we limit our search domain to training trials with the same inter-onset-interval and standard tone as the misjudgement trial. In the next step, we search for correlations between the perception and various oddball tone properties such as its 1) onset (time step at which the oddball begins, *T*_init_), 2) duration (Δ*T*), and 3) ending time point (time point at which the oddball ends, *T_fin_*). To this end, we calculated the information shared between the perception and oddball tone properties (see Methods for a detailed explanation of information computation procedures). Fig. 6A shows the information shared between the perception (decision state *S*_decision_) of the Brains and 1) oddball ending time (shown in grey), 2) oddball onset (shown in blue), and 3) oddball duration for each inter-onset-interval and standard tone. These results show that the oddball ending time point is a better predictor of the perception than the oddball tone onset or its duration. Note also that the information shared between the perception and the oddball ending time point remains consistent across all IOI and standard tones, whereas shared information between perception and oddball duration, and perception and onset decrease monotonically as IOI and standard tones increase. Building on these results, we propose the following alternative hypothesis: during misjudgement of an early or late oddball, a Brain goes through a state sequence that is reminiscent of experiences it had during trials with the same IOI and standard tone, and with on-time oddballs that end at the same time point as the early or late oddball (an example scenario is shown Fig. 6B).

**Fig. 6.**
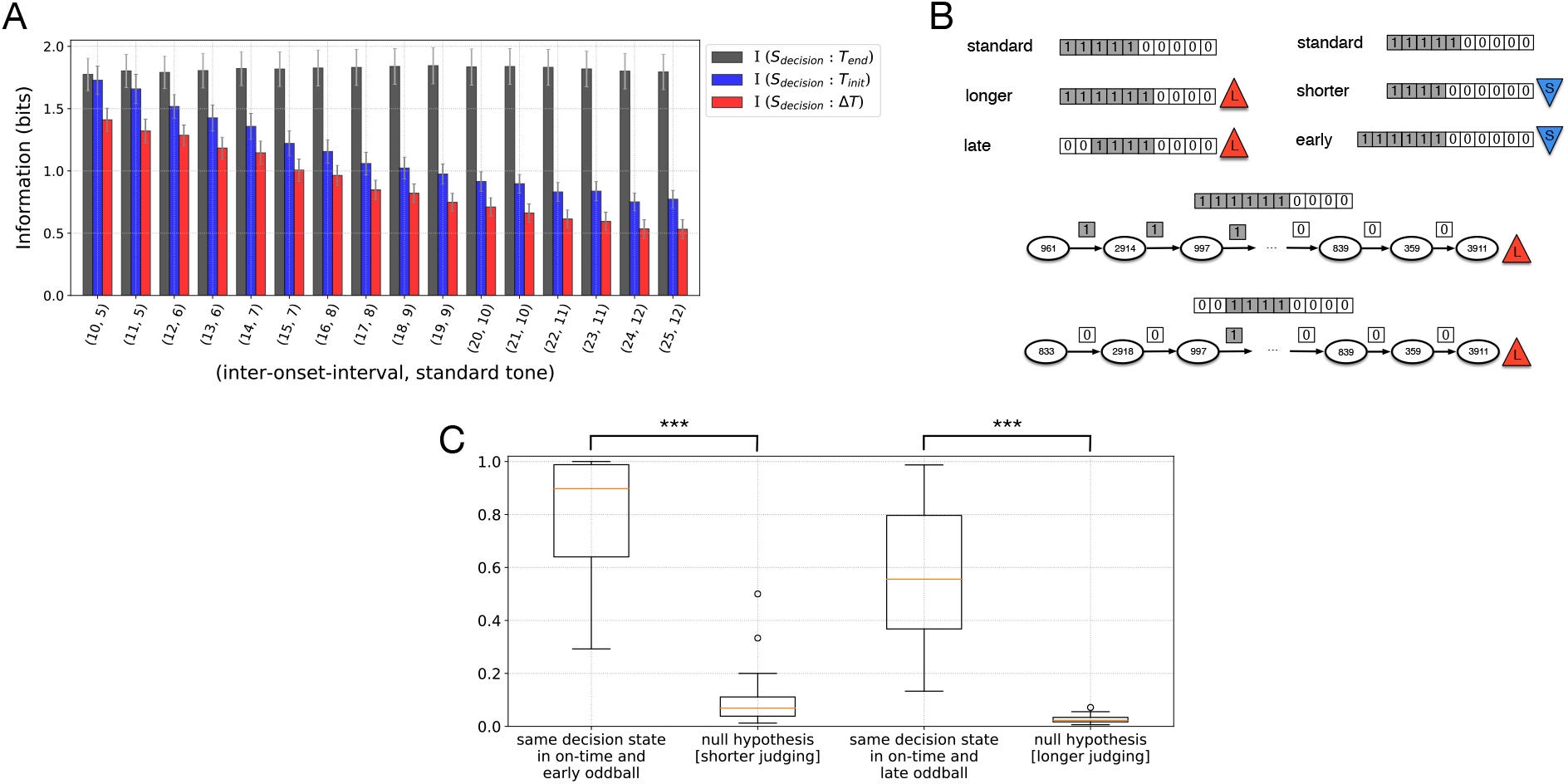
(A) The mutual information between perception, i.e., the decision state of the Brain, and 1) the oddball tone ending time step (shown in black), 2) the oddball tone duration (shown in red), 3) the oddball tone onset (shown in blue), and their 95% confidence intervals. (B) Sequence of inputs for a standard tone, an on-time longer oddball tone that is correctly judged as longer, and a shorter late oddball tone that is misjudged as longer. Sequence of inputs for a standard tone, an on-time shorter oddball tone that is correctly judged as shorter, and a longer early oddball tone that is misjudged as shorter. Sequences of Brain states along with input sequences for on-time longer oddballs and shorter late oddballs.(C) The fraction of misperceived out-of-time oddball tones that resulted from having the same perception in on-time and out-of-time stimuli with the same oddball end points (left data point), compared to the null hypothesis; likelihood that Brains misjudgements were to be issued from any one of states from set of “shorter-judging” or “longer-judging” states (middle and right data point, respectively).

In order to test this alternative hypothesis, we perform another test to measure how often perception in misjudgement of early or late oddballs is identical to perceptions in similar training trials. Consider for example a trial with IOI=10, standard tone=5 with a late oddball tone (onset=2) that is shorter than the standard tone (duration=4) as shown in Fig. 6B. When a Brain misjudges this oddball as “longer” (with *S*_decision_ = 3911 as shown in Fig. 6B), we search for instances in the set of training trials (with on-time oddball) with IOI=10 and standard tone=5, where that Brain issued a correct “longer” decision for an oddball that ended at the same time point as the late shorter oddball (as shown in Fig. 6B). The same analysis can be performed for misjudgement of early oddballs that are longer than the standard tone (Fig. 6B). We count the number of such instances for each Brain and divide the result by the total number of its misjudgements of out-of-rhythm oddball tones. Fig. 6C (left data point) shows the result of this analysis for all 50 Brains. This result shows that in the vast majority of the cases (with median of 69.5% of the cases), the misjudged out-of-rhythm oddball and on-time oddballs that end at the same time point are *perceived* the same. In other words, the misjudgement is due to Brains paying less attention to the onset of the tone, meaning the onset of the oddball does not affect the ultimate state from which the decision is issued. The middle and left data points show the probabilities calculated from equation 1 described in the null hypothesis, that measure how likely it is for a Brain to, by chance, end up in any of the “shorter-judging” or “longer-judging” state at decision time. Our statistical analysis shows that having the same decision state in out-of-rhythm oddball and on-time oddballs (with constraints explained above) are significantly more likely than being in “shorter-judging” (median=0.695 vs. median=0.069, Mann Whitney *U* = 2494.0, *n* = 50, *p* = 5.03 × 10^−18^ one-tailed) or “longer-judging” state at decision time (median=0.695 vs. median=0.023, Mann Whitney *U* = 2500.0, *n* = 50, *p* = 3.51 × 10^−18^ one-tailed), therefore, we reject the null hypothesis in favour of the alternative hypothesis.

Based on these findings, we conclude that during misjudgements of early or late oddball tones, Markov Brains pay more attention to the end point of the oddball and less attention to the oddball duration, or it onset. This is presumably because during evolution tones are always rhythmic and Brains that entrain to the rhythm expect the oddball to be on-time. As a result, Brains pay more attention to when the oddball ends which is a more informative component of the stimuli than its onset which, during evolution, had no variation and hence, no uncertainty.

#### Experience of early or late oddball is similar to adapting entrainment to phase change

Here we investigate the entire sequence of Brain states (Brain experiences of the stimuli) for those instances we found in previous section, in which the perception of the Brain in misjudgement of early/late oddballs was the same as perception of shorter/longer on-time tones with the same end point as an out-of-time oddball. In order to compare two experiences, we use two different measures (experience comparison is a form of representational similarity analysis, see for example (45, 46)). First, we find the longest common sub-sequence that includes the decision state. In other words, we start from the decision state in on-time and out-of-time sequences (note that the decision state is the same in both sequences), trace back the transitions in sequences and count the number of states that are identical in both sequences until the first mismatch occurs. The length of the identical portion of the two sequences is then normalised by the total length of one sequence (recall that the length of both sequences are the same) to lie in the range (0,1], we term this normalized length of the identical portion of experiences the *similarity depth*, since it measures how deeply the on-time and out-of-time oddball experiences are identical. We note that because the perception of the tone is the same in these trials, the similarity depth must be greater than zero. Second, we use the Jaccard index, that measures the overall similarity of sequences by comparing states at same positions in the two sequences.

Fig. 7A shows the distributions of similarity depth and total similarity of experiences. Fig. 7B shows the distribution of the difference between the similarity depth and total similarity. The difference between the two measures is zero in 91.5% of the cases which implies that the experiences are almost always entirely different up to the point where they become identical. We observe a wide variety in these similarity measures which shows that Brains do not traverse the exact same trajectory they did during an on-time trial; rather the early or late oddball initially throws the Brain out of this trajectory but later the Brain returns to states it experienced during an on-time oddball with the same end point. In other words, the onset of the out-of-time oddball is noticed, however, since the Brains are entrained to the rhythm and expect the oddball to be on-time their computations of duration relies more on their expectation than the actual start point of the oddball. This mechanism is reminiscent of adapting to phase changes in entrainment to rhythmic stimuli.

**Fig. 7.**
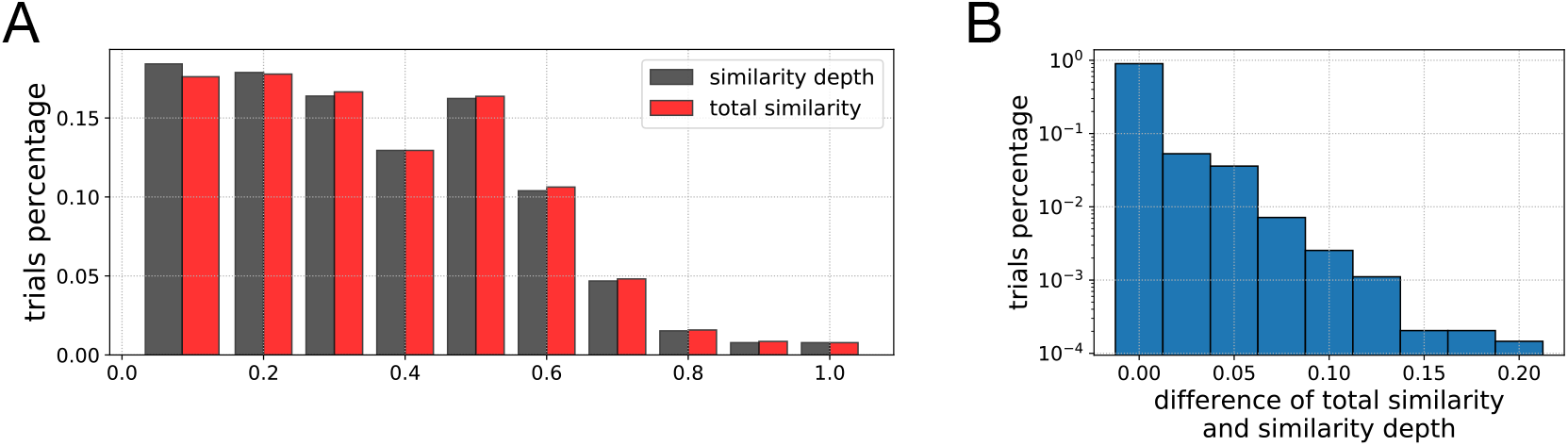
(A) Distribution of similarity depth of experiences (sequences of states) of on-time and early/late oddball tones in trials in which onset does not change the perception of the tone in Markov Brains. Similarity depth one implies that the experiences are identical throughout the tone perception. (B) The distribution of the difference between the total similarity and similarity depth in each trial.

## Discussion

This study was aimed at elucidating the neural (mechanistic) underpinnings of perception, by evolving digital Brains that perform duration judgements of tones that were presented in a rhythmic sequence, and that were later subjected to out-of-rhythm oddball tones to quantify distortions in duration judgement that occur as a response to the onset manipulation. We found that evolved Markov Brains display a capacity to discriminate tone length that is remarkably similar to people’s ability to distinguish changes (quantified by Weber’s Law) to the extent that the observed relative JND of Markov Brains was in the same range (6-10%) as in some of the experiments in (24, 40). Furthermore, evolved Markov Brains exhibit a systematic distortion in perceived event duration of out-of-rhythm oddball tones that is also similar to what was observed in a human subjects study previously conducted by one of the authors. But while the conclusion of (24) was that the experiments supported the dynamic attending theory (DAT) of attentional entrainment (which, we recall, posits that entrainment creates peaks of attention that coincide with the start of each tone) we here find instead that Markov Brains pay attention to the *end of the signal*, and pay less attention to the onset.

From the point of view of Bayesian inference (47), a model of cognition that focuses attention on those parts of the signal that carry most of the uncertainty (the end of the stimulus) makes eminent sense. After all, Brains that have experienced only on-time stimuli should take the rhythmic nature of stimuli for granted: there is no need to pay attention to predictable stimuli. In fact, this view of cognition is fully consistent with the theory that perception is an inference problem that focuses on prediction-error reduction (48), as well as the Hierarchical Temporal Memory (HTM) model of neo-cortical computation (49), which is based on the idea that brains are prediction machines. This model of attention differs from common models of visual processing and attention such as visual and auditory saliency (50, 51), because in those models only the contrast of the stimulus with the background is considered for saliency, not the value of the information it contains. The model is consistent, however, with neurophysical models in which temporal anticipation improves perception but does *not* affect the spontaneous firing rate (36, 37), which is associated with attention in visual processing (52).

The present work suggests a model of cognition where the stimulus not only entrains the cognitive apparatus, but conditions the brain to expect only a small subset of possible future states. From this point of view, any temporal history of stimuli leads to predictions that, for the most time, will come to pass unless the environment has changed in a way that necessitates further attention. In particular, our findings suggest that both DAT and SET are incomplete models of time perception where DAT unduly emphasises attention peaks at the beginning of each tone in the sequence, while SET uses the onset and the end of the tone to start and stop a clock, contrary to our (admittedly digital) evidence.

The results presented here open up a number of different questions and avenues for future exploration. Can the theory of dynamical entrainment we present here be meaningfully tested in human experiments, by focusing on those predictions that distinguish it from established theories such as SET and DAT? Does this theory also explain observations in different sensory modalities such as vision? A program in which empirical studies using human subjects coupled with sophisticated digital experimentation might provide an answer, and open up avenues for a detailed mechanistic understanding of the complexities of perception. Ultimately, this opens up the possibility of explaining phenomenological concepts such as attention, perception, and memory in terms of state-space dynamics of cortical networks.

## Methods

The use of mathematical and computational methods for the study of behaviour is growing, especially due to the unprecedented increase in our computational power (53). Computational methods in particular enable us to perform a large number of “experiments” *in silico*, with parameters varying in a wide range, in a reasonably short time. Such experiments allow us to explore parameter space more broadly and to make predictions about conditions that have not been tested before and, more importantly, are currently beyond the reach of our empirical power. Naturally, for such computational experiments to have any explanatory power, they must be validated thoroughly with behavioural data.

In this work, we use an agent-based model in which agents are controlled by artificial neural networks (ANNs) that differ in many important aspects from the more common ANN method. Because the logic of these networks is determined by logic gates with the Markov property we refer to these neural networks as Markov Brains (54). Below, we describe the structure, function, and encoding of Markov Brains, but see (34) for a full description of their properties and how they are implemented. Markov Brains have been shown to be well-suited for modelling different types of behaviour observed in nature, from simple elements of cognition such as motion detection (55) and active categorical perception (41, 56), to swarming in predator-prey interactions (57), foraging (58), and decision-making strategies in humans (59).

### Markov Brains

Markov Brains are networks of variables connected via probabilistic or deterministic logic gates with the Markov property. While we often term these variables “neurons”, the state of the variable is more akin to a binary firing *rate*, that is, each neuron is a binary random variable (i.e., a bit) that may take two values: 0 for quiescent and 1 for firing. Fig. 8A shows a schematic of a simple Brain consisting of 12 neurons (labelled as 0-11) at two subsequent time points *t* and *t* + 1. The state of neurons in this example are updated via two logic gates. Fig. 8B shows a gate that takes inputs from neurons 0, 2, and 6 and writes the output into neurons 6 and 7. This logic gate produces output states of neurons 6 and 7 at time *t* + 1 given input states at time *t*. Each gate is defined by a probabilistic logic table in which the probability of each output pattern for a given input is specified. For example, in the probability table shown in Fig. 8C, *p*_52_ specifies the probability of obtaining output state (*N*_6_,*N*_7_) = (1,0) (a state with decimal representation ‘2’) given input states (*N*_0_,*N*_2_,*N*_6_) = (1,0,1) (decimal translation ‘5’), that is,

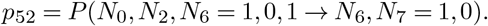

**Fig. 8.**
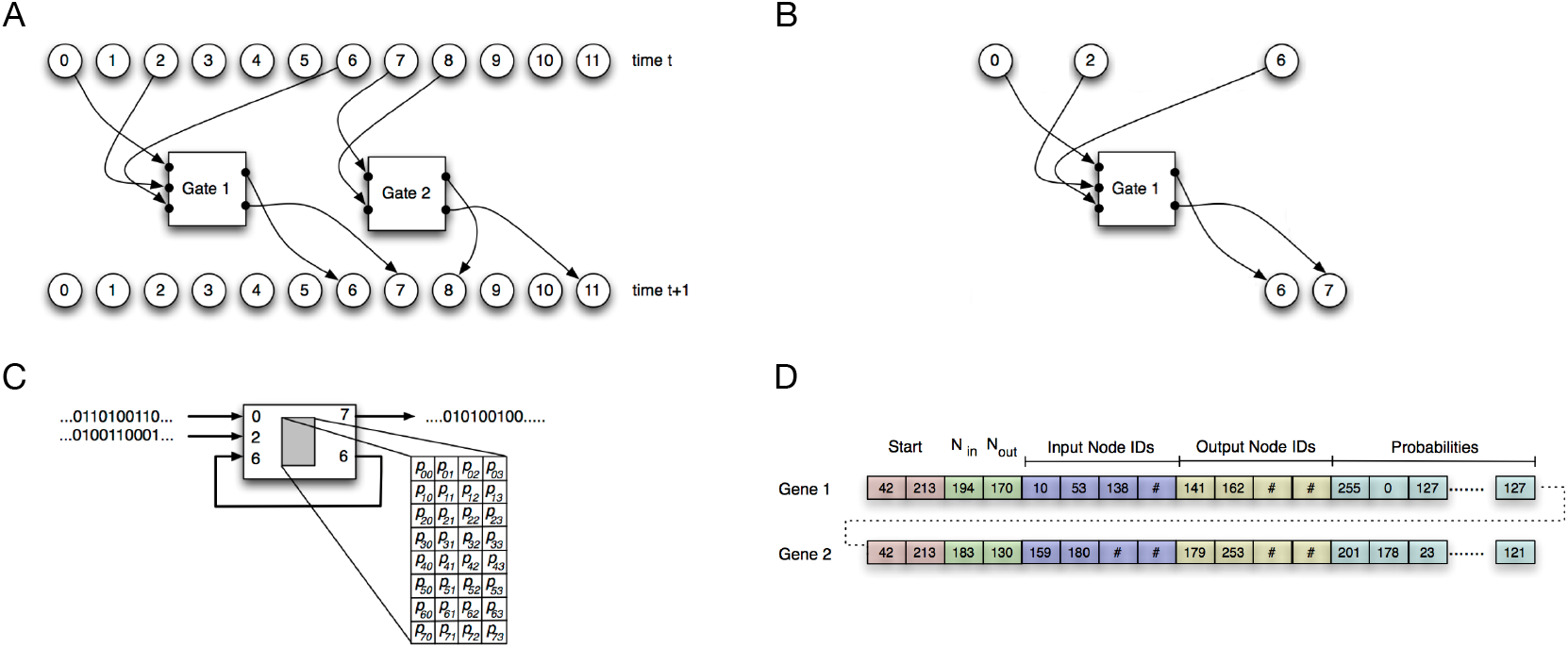
(A) A simple Markov Brain with 12 neurons and two logic gates at two consecutive time steps *t* and *t* + 1. (B) Gate 1 of (A) with 3 input neurons and 2 output neurons. (C) Underlying probabilistic logic table of gate 1. (D) Markov Network Brains are encoded using sequences of numbers (bytes) that serve as agent’s genome. This example shows two genes that specify the logic gates shown in (A), so that, for example, the byte value ‘194’ that specifies the number of inputs *N*_in_ to gate 1 translates to ‘3’ (the number of inputs for that gate).

Since this gate takes 3 inputs, 2^3^ possible inputs can occur, which are shown in eight rows. Similarly, this probabilistic table has four columns, one for each of the 2^2^ possible outputs. The sum of the probabilities in each row must equal 1: ∑_*j*_ *p_ij_* = 1. When using deterministic logic gates (such as in this study), all the conditional probabilities *p_ij_* are zeros or ones. In general, Markov Brains can contain an arbitrary number of gates, with any possible connection patterns, and arbitrary probability values in logic tables (34). As is clear from this example, we do not implement the update of the Brain state using probabilities that are conditional on the environmental state 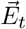; rather, we update the joint state 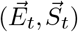.

In Markov Brains, a subset of the neurons is designated as sensory neurons that receive inputs from the environment. Similarly, another subset of neurons serves as actuator neurons (or decision neurons) that enable agents to take actions in their environment. In principle, an optimal Brain is designed in such a manner that a particular sequence of inputs (a time series of environmental states 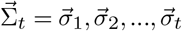) leads to a Brain state 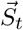 that triggers the optimal response in that environment. Rather than using an optimisation procedure that maximises an agent’s performance over the probabilities 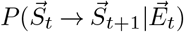, we use an evolutionary process in which a Brain’s entire network is encoded in a genome (60) and optimisation occurs through the evolution of a population of such genomes using a “Genetic Algorithm” (GA, see for example (61)). In particular, each gene specifies a gate’s connectivity and its underlying logic as shown in Fig. 8D. This evolutionary approach is explained in more detail in the following section.

### Evolution of Markov Brains

Markov Brains can evolve to perform a variety of tasks representing different types of behaviours observed in nature. Selecting for any desirable task leads to the evolution of network connections and logic-gate properties that enable the agents to succeed in their environment. Each genome is a sequence of numbers ranging between 0-255 (bytes) that represent a set of genes that encode the logic and connectivity of the network. The arbitrary pair of bytes 〈42,213〉 represents the “start codon” for each gate (Fig. 8D), while the downstream loci instruct the compiler how to construct the network, by encoding how many inputs and outputs define each logic gate, where the inputs come from (that is, which neuron or neurons), and where it writes to. In this manner, by “expressing” each gene, the network is fully determined via the connections between neurons and the logic those connections entail. Once a Brain is constructed, it is implanted in an agent whose performance is evaluated in an artificial environment that selects for the task. Those agents that perform best are rewarded with a differential fitness advantage. As these genomes are subject to mutation, heritability, and selection, they evolve in a purely Darwinian fashion (albeit asexually). The Genetic Algorithm specification details are shown in Table 2.

**Table 2.**
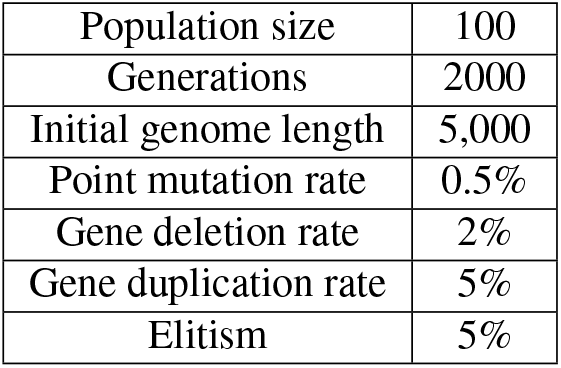
Genetic Algorithm configuration. We evolved 50 populations of Markov Brains for 2,000 generations with point mutations, deletions, and insertions. We used roulette wheel selection, with 5% elitism, and with no cross-over or immigration.

|  |  |
| --- | --- |
| Population size | 100 |
| Generations | 2000 |
| Initial genome length | 5,000 |
| Point mutation rate | 0.5% |
| Gene deletion rate | 2% |
| Gene duplication rate | 5% |
| Elitism | 5% |

The population of Markov Brains evolves to judge the duration of an oddball tone (“longer” or “shorter”) in multiple trials with different IOIs and oddball durations. The full set of all (IOI, standard tone), possible oddball tone durations, and the total number of trials for each pair of (IOI, standard tone) used in the evolution is shown in table 3. All told, there are 1,472 possible trials. However, agents are only evaluated on a subset of trials in every generation. This sampling increases the evolution efficiency (62), and helps to avoid overfitting and enhances generalisation of learning (63). In each generation, we randomly pick 22 trials from each (IOI, standard tone) pair (each row in Table 3) to form the evaluation subset: 11 trials with a longer oddball, and 11 trials with a shorter oddball, so as to prevent biasing Brains toward one response or the other. All agents of the population are then evaluated in that same subset of trials, which is 352 trials.

**Table 3.** Complete set of all inter-onset-intervals, standard tones, and oddball durations used for the evolution of duration judgement. Oddballs can occur in either of the 5th, 6th, 7th, or 8th position in the rhythmic sequence. Also, oddball durations are always either shorter or longer than the standard tone. The total number of trials for each pair 〈ioi, tone〉 is four times the IOI minus 2 (excluding oddball duration=standard tone, oddball duration=IOI), because the oddball can appear in four different positions within the rhythmic sequence.

| (Inter-onset-interval, Standard tone) | Oddball tone durations | Total number of possible trials | Number of evaluation trials |
| --- | --- | --- | --- |
| (10, 5) | {1, 2, 3, 4}, {6, 7, 8, 9} | 32 | 22 |
| (11, 5) | {1, 2, ..., 4}, {6, ..., 10} | 36 | 22 |
| (12, 6) | {1, 2, ..., 5}, {7, ..., 11} | 40 | 22 |
| (13, 6) | {1, 2, ..., 5}, {7, ..., 12} | 44 | 22 |
| (14, 7) | {1, 2, ..., 6}, {8, ..., 13} | 48 | 22 |
| (15, 7) | {1, 2, ..., 6}, {8, ..., 14} | 52 | 22 |
| (16, 8) | {1, 2, ..., 7}, {9, ..., 15} | 56 | 22 |
| (17, 8) | {1, 2, ..., 7}, {9, ..., 16} | 60 | 22 |
| (18, 9) | {1, 2, ..., 8}, {10, ..., 17} | 64 | 22 |
| (19, 9) | {1, 2, ..., 8}, {10, ..., 18} | 68 | 22 |
| (20, 10) | {1, 2, ..., 9}, {11, ..., 19} | 72 | 22 |
| (21, 10) | {1, 2, ..., 9}, {11, ..., 20} | 76 | 22 |
| (22, 11) | {1, 2, ..., 10}, {12, ..., 21} | 80 | 22 |
| (23, 11) | {1, 2, ..., 10}, {12, ..., 22} | 84 | 22 |
| (24, 12) | {1, 2, ..., 11}, {13, ..., 23} | 88 | 22 |
| (25, 12) | {1, 2, ..., 11}, {13, ..., 24} | 92 | 22 |

### Experimental Setup

The Brains we evolve can have up to 16 neurons, of which one serves as the sensory neuron, and one delivers the decision (the “actuator” neuron). The remaining 14 neurons can be used for computation and signal transduction, but how many of them are actually used is determined by evolution. The population of Markov Brains evolves to judge the duration of a deviant tone (oddball) within a rhythmic sequence of otherwise identical tones, similar to experiments in (24) (see Fig. 9). In each trial, agents listen to a sequence of nine tones with a constant inter-onset-interval (IOI). An oddball is embedded within this sequence that is either shorter or longer in duration compared to the other eight tones (standard tones). Markov Brains sense the stimulus in one of their neurons (here, neuron 0, see Fig. 9). Agents must decide whether the oddball stimulus is longer or shorter than the standard tones. The agent is rewarded for correct duration judgements and does not gain any reward or incur a penalty for incorrect judgements. One neuron (neuron 15) in the Markov Brain is designated for delivering the decision (“longer” or “shorter”).

**Fig. 9.**
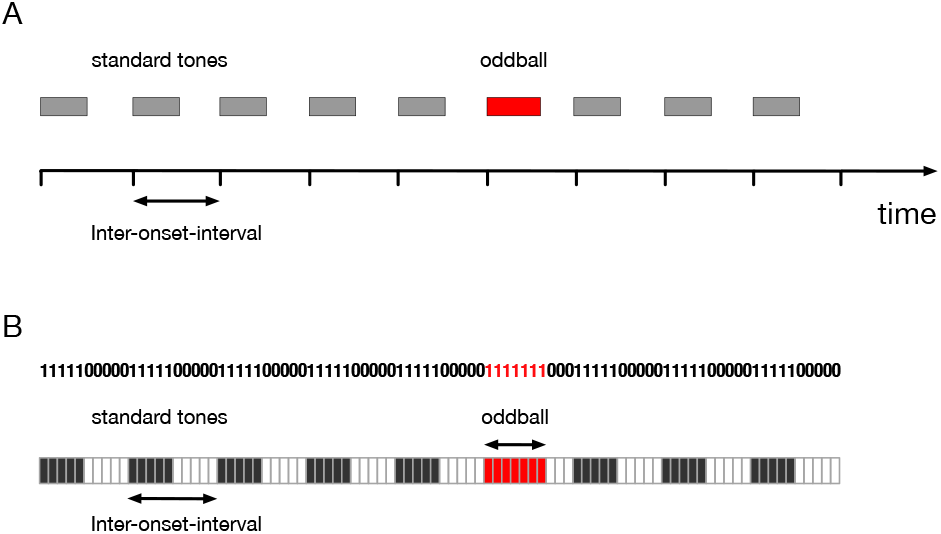
(A) A schematic of auditory oddball paradigm in which an oddball tone is placed within a rhythmic sequence of tones, i.e., standard tones. Standard tones are shown as grey blocks and the oddball tone is shown as a red block. (B) The oddball auditory paradigm, which is converted to a sequence of binary values, shown as sensed by the input neuron of a Markov Brain. When a stimulus is present, a sequence of ‘1’s (shown by black blocks) is supplied to the sensory neuron while during silence, a sequence of ‘0’ is fed to the sensory neuron. Each block shows one time step of the sequence experienced by the Brain.

For the purpose of fitness evaluation, agents are evaluated in several trials with different inter-onset-intervals (IOIs), dif-ferent standard tones, a wide range of oddball durations, and with oddballs placed in different positions in the sequence. Standard tones range from 5 time steps to 12 time steps. The IOI is approximately twice the standard tone, and ranges from 10 to 25. Oddball durations can take any value from the shortest possible duration (1 time step) all the way to IOI minus 1 to avoid interfering with the next tone. During evolution, agents are not evaluated with oddball tones with the same duration as the standard tone since it is not shorter or longer than the standard tone. Oddballs can occur in either 5th, 6th, 7th, or 8th position, exactly as in the protocol of (24). Our standard tones would be comparable in duration to those used in (24) if a digital time step is represented by a physical signal with about 70msec duration.

The set of all IOIs, standard tones, possible oddball-tone durations, and the total number of trials for each pair of (IOI, tone) is given in Table 3. All agents of the population are then evaluated in that same subset of trials, half of which with a longer oddball and the other half with shorter oddball, to avoid creating a bias in the agents’ judgements. This subset of randomly picked trials consists of 512 trials (out of a total 2,852 trials): 22 trials for each (inter-onset-interval, standard tone) (see Table 3).

### Discrete time in Markov Brains

The logic of Markov Brains is implemented by probabilistic or deterministic logic gates that update the Brain states from time *t* to time *t* +1, which implies that time is discretised not only for Brain updates, but for the environment as well. Whether or not the brain perceives time discretely or continuously is a hotly debated topic (64), but for common visual tasks such as motion perception (65) discrete sampling of visual scenes can be assumed. For Markov Brains, the discreteness of time is a computational necessity. Because no other states (besides the neurons at time *t*) influence a Brain’s state at time *t* + 1, the gates possess the Markov property (hence the name of the networks). Note that even though the Markov property is usually referred to as the “memoryless” property of stochastic systems, this does not imply that Markov Brains cannot have memory. Rather, memory can be explicitly implemented by gates whose outputs are written into the inputs of other gates, or even the same gates, i.e., to itself (41, 54).

### Markov Brains as finite state machines

Because the Brains we evolve are deterministic, they effectively represent a deterministic finite-state automaton (DFA). There is considerable literature covering the mathematics of DFAs (see, for example (66)), but very little is applicable to the automata we evolve here. For example, realistic evolved automata are unlikely to have absorbing states, their stationary distributions are irrelevant, and they may be both cyclic and acyclic.

We define the state of a Markov Brain as the vector of states of all neurons except the sensory ones (35, 43, 67): 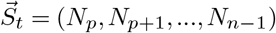, where *N_i_* is the state of the *i^th^* neuron, *p* is the number of sensory (or peripheral) neurons, (*N*_0_, *N*_1_,…, *N*_*p*−1_) is the state vector of sensory neurons, and *n* is the total number of neurons. We abbreviate the Brain-state using the decimal translation of the state vector as:

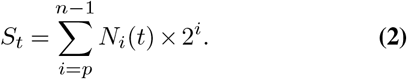

The Brain state can be thought of as a snapshot of the entire Brain that contains information about the activity (firing rate) of all neurons at that particular point in time. Markov Brains go through discrete states as the agent it controls behaves, reminiscent of what has been observed in monkeys performing a localisation task (43). In our experimental setup, Markov Brains have 16 neurons in total, so *n* =15. One of the neurons senses the stimulus, i.e. *p* = 1, so equation [2] can be written as 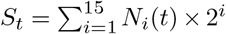 which means the Brain can be in at most 2^15^ = 32,768 different states. We also denote the sensory input at time *t* as 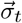, and define the sequence of sensory inputs from time *t*_0_ to 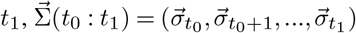.

The initial Brain state is always 0 since all neurons are quiescent at the outset. State-to-state transitions of an evolved Brain can be represented (or explained) as a mapping of the state of the Brain and the sensory input to the future state of the Brain. Formally, the set of all transitions of the Brain over all *visited* states in trials (states that Brains have taken on in those trials) can be viewed as a function 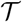 that takes the current state of the Brain *S_t_* as well as the sensory input 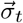 (in our experimental setup it is just one bit) as the input, and returns the future state of the Brain as the output, *S*_*t*+1_:

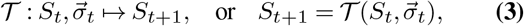

We restrict the domain of variable *S_t_* to those Brain states that actually occur during training (i.e., evolution) or test trials (early/late oddball tones). This function can be illustrated as a directed graph in which Brain states are represented by nodes (labelled by the decimal translation of the Brain state, see Eq. [2]) and edges represent transitions that are labelled with the stimulus that drives those transitions, 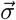 (see (34) for a more detailed exposition of state-to-state diagrams).

### Attention, experience, and perception in Markov Brains

We describe Markov Brains in terms of functions that take 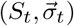 as the input and return *S*_*t*+1_ as the output.

#### Definition 1

If the Brain transitions from a particular state *S_t_* to the same state *S*_*t*+1_ for all possible values of 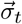 we say: the Brain *does not pay attention* to sensory input 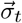 in state *S_t_*.

Note that it is possible that the Brain does not pay attention to *parts* of the sensory input 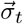 when the transition from *S_t_* to *S*_*t*+1_ occurs independently of specific components of vector 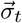. We emphasise that when the Brain does not pay attention to a sensory input in one transition, it does not imply that the stimulus is not sensed. Rather, it implies that even though sensed, the value does not affect the Brain’s computation when in state *S_t_*. It is crucial here that this definition of attention to a stimulus depends not only on the stimulus itself but also on the context in which it is sensed—this context is represented by the state *S_t_* the Brain has reached. Because the Brain has reached the state *S_t_* as a consequence of the temporal sequence of states traversed, this context is in fact historical. Also, note that the Brain state encompasses the actuator neuron (decision neuron), therefore, “not paying attention” is reflected in an agent’s behaviour as well as the Brain’s computations on sensory information. In a sense, the definition implies that an event that the Brain does not pay attention to should not alter its *experience* of the world, a concept that we will now define.

#### Definition 2

We define the Brain’s experience of the environment (which is sensed as a sequence of sensory inputs 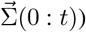 as the sequence of Brain states it traverses, i.e., as 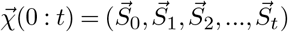.

This definition implies that the experiences of different individual Brains can be different when encountering the exact same sensory sequence, hence, experience is subjective (68, 69). Furthermore, an agent may have experiences in which it does not take any actions on its environment (does not make any physical changes to itself or the world). Thus, dreaming or thinking are instances of such experiences in humans (68–70). However, if the agent takes any actions in its environment, those actions become part of the experience by definition. For example, in our experimental setup Brains can only “take an action” in one particular time step of the trials. As a result, a sequence of states that excludes that time step is still an experience, but does not involve any actions from the agent. It is also crucial to understand that the experience of the environment that is represented within Brain states is not just a naive projection of the world on the Brain, but rather contains integrated information about the relevant aspects of the environment (cues), while ignoring unimportant details (noise). In a very real sense, a Brain separates signal from noise; information from entropy (71).

In general, two different input sequences 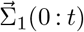 and 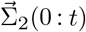 will result in the Brain having two different experiences 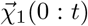 and 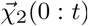, but not necessarily. If experiences 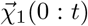 and 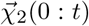 are exactly the same, it means that (according to Definitions 1 and 2) the Brain does not pay attention to inputs during those transitions in which 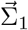 and 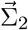 are different. While in Definition 1 we only considered the Brain’s transition at one time step, we can also look at the sequence of *future* Brain states, to discover how sensory inputs affect the Brain’s computations and transitions multiple time steps after the input is sensed. Now, consider two input sequences 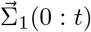 and 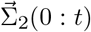 that differ in time steps (0: *t*′), where *t*′ < *t*. Also, suppose 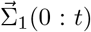 and 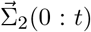 result in two different experiences 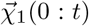 and 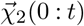. The effect of sub-sequence 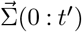 can be gauged by how different experiences 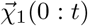 and 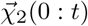 are as a result. For example, if two input sequences 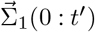 and 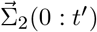 (during time interval 0: *t*′ where they are different) throw the Brain into two different regions in state space and therefore give rise to completely different experiences, then those inputs disturb experiences substantially. If, by contrast, 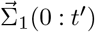 and 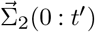 only result in different experiences temporarily (for example, during 0: *t*′) while 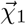 and 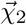 become similar or identical later, then the differences in inputs is less disruptive to the Brain’s experience. In particular, if the experiences have identical states at decision time *t_d_* (assuming that *t_d_* ∈ [0: *t*]), the differences in sensory inputs impact experiences 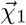 and 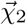 even less. We emphasise that the Brain state at the point of decision is key, because at this time point in the trial, the state of the Brain specifies the Brain’s judgement, and more importantly, represents the path traversed in state space to reach this state. Consequently, we use the Brain state at decision time to define what it means to “perceive” a sensory input sequence.

#### Definition 3

If a Brain encounters two different input sequences 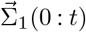 and 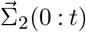, yet ends up in the *same* state *S_t_* at decision time *t* in both cases, we say that the Brain *had the same perception* of sensory sequences 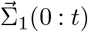 and 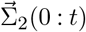.

By this definition, “having the same perception” is a superset of “having the exact same experience” when encountering two different sensory sequences. As discussed earlier, if the Brain has the exact same experience when exposed to two different input sequences, it clearly does not pay attention to the sub-sequence of the inputs that is not common between the two input sequences. In the same vein, how similar the experiences are for two different input sequences correlates with how little the Brain pays attention to those parts of input sequences that are not the same. This correlation captures the idea that there are different levels of “not paying attention” to a phenomenon in the environment. At the same time, it becomes clear that events that evoke the same perception (and thus similar experiences) must overlap in the significant parts of the sensory input. In this manner, the state of the Brain— being specific to the path in state space that leads to it—can encode “involuntary memory”, in the same way as Marcel Proust’s memories of the past (72) are triggered by the taste of a Madeleine dipped in Linden tea.

### Information shared between perception and the oddball tone

Here we describe the procedures used to calculate the information shared between perception, (the Brain state at decision time-step), and the different oddball tone properties such as its duration, onset, and ending time-step. Markov Brains are tested against oddball tones varying in durations as well as different onsets with respect to the rhythm of the sequence. For each individual Brain we create an ensemble of trials with the same inter-onset-interval and standard tone, in which oddball tones differ in duration, onset, or both. We can calculate the information shared between the perception of each individual Brain and oddball properties for a given interonset-interval and standard tone using the standard Shannon information (73)

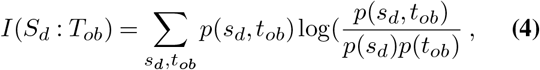

where *S_d_* denotes the Brain state at decision time (which we defined as perception) and *T_ob_* denotes oddball properties, for example the oddball duration. The shared information between the perception and the oddball properties (duration, onset, and ending time-step) captures the correlation between the perception of the Brain and each of the oddball properties. It is noteworthy that perception occurs after the oddball tone has arrived and terminated. Thus, the information Eq. (4) measures how well each of the oddball tone properties can predict how the Brain perceives the tone.

## Data accessibility

All source code used to conduct this study, as well as post-processing scripts, are available at the repository https://github.com/AliTehrani/Attentional-Entrainment

## Author contributions

A.T., J.D.M., and C.A. designed research; A.T. performed research and analysed data; and A.T., J.D.M. and C.A. wrote the paper.

## Competing Interests

We declare we have no competing interests.

## Funding

This material is based in part upon work supported by the National Science Foundation under Cooperative Agreement No. DBI-0939454.

## Acknowledgements

This work was supported in part by Michigan State University through computational resources provided by the Institute for Cyber-Enabled Research. We thank members of the Adami Lab, Hintze Lab, and Dr. Taosheng Liu for helpful discussions.

## Supplementary Information

**Supplementary Figure 1.**
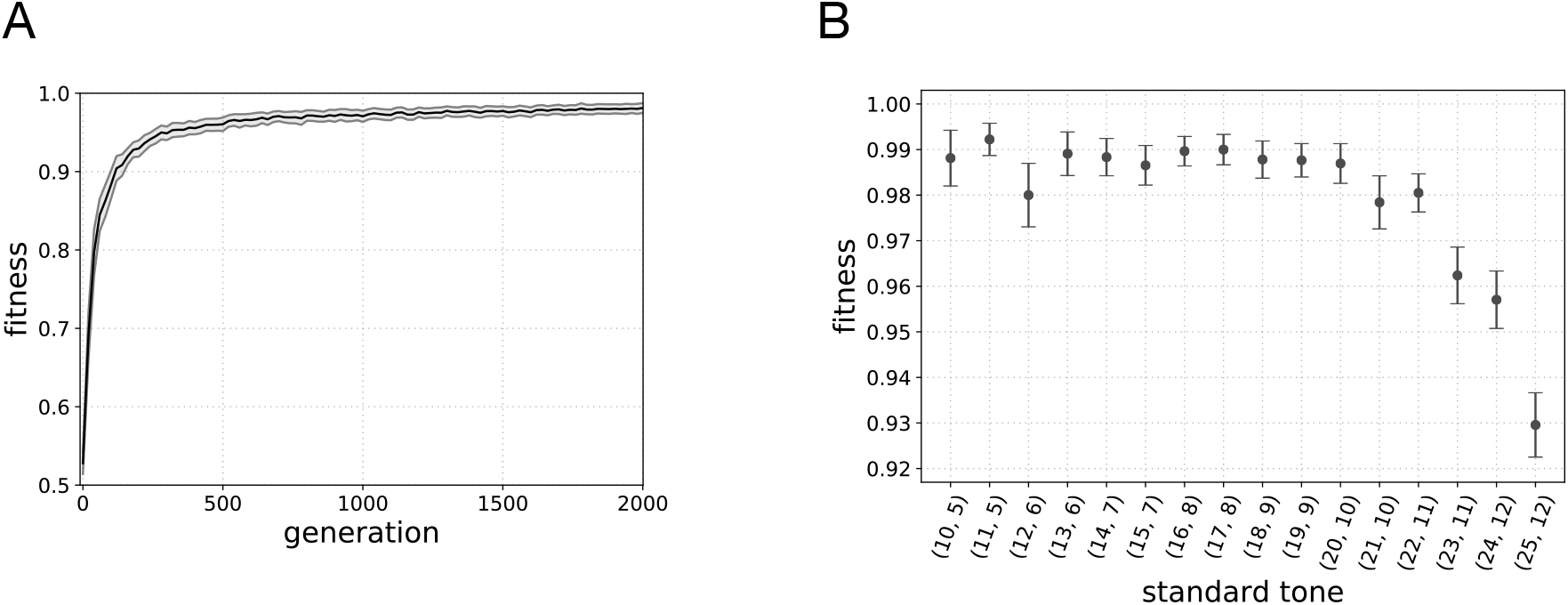
(A) Mean fitness across all 50 lineages and 95% confidence interval as a function of generation shown every 20 generations. (B) Mean fitness (and 95% intervals) of best agents picked from each of the 50 populations after 2000 generations as a function of inter-onset-interval, standard tone.

**Supplementary Figure 2.**
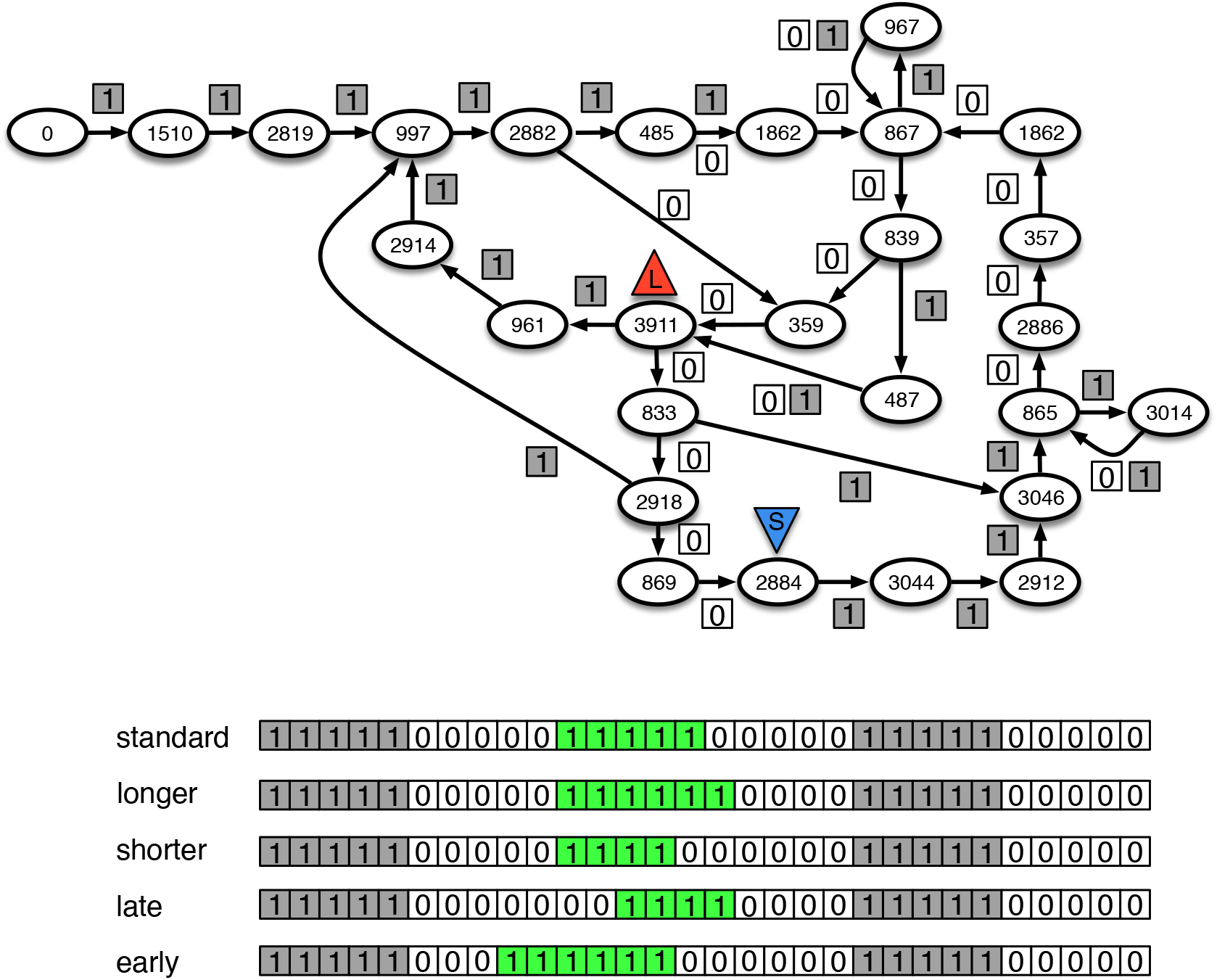
State-to-state transition diagram of a Markov Brain for IOI=10, standard tone=5, oddball tones=4 and 6, and onset of oddball tones can be 2 time steps early and 2 time step late.

### Fitness landscape structure and historical contingencies result in Markov Brains using smaller regions of state space in trials with longer IOIs

In the main text we described that the judgement accuracy deteriorates as the IOI (and therefore tone lengths) increases. More specifically, even though the relative JND values remain in the same range for different IOI and standard tones (see Fig. 2B in the main text), PSE values start to deviate from the standard tone leading to higher values of “constant errors” (CE) that is, the difference between PSE and POE (see Fig. 2C in the main text). Here, we show that 1) deviations of PSEs in longer IOIs result from the fitness landscape structure and historical contingencies (see for example (1, 2)), and 2) the mechanistic basis of these deviations is associated with the size of the state-space Markov Brains use to encode stimuli characteristics.

As discussed before, Markov Brains display periodic firing patterns in response to rhythmic stimuli. These periodic patterns result in the formation of loops in their state transitions. This is the dominant mechanism by which Brains evolve to entrain to rhythmic stimuli, and encode temporal characteristics of the stimuli (i.e., rhythm and standard tone’s duration). The distribution of the period of these periodic firing patterns, that is, the lengths of the loops in state transition diagrams is shown here again in Supplementary Fig. 3A. Since the first four standard tones are provided so that Brains entrain to the rhythm, we measured the period of state transitions after the first four intervals, without an oddball tone. We also measured the number of distinct states each Brain visits during these periodic state transitions. Supplementary Fig. 3B shows the distribution of number of distinct states in traversing loops during entrainment for 50 evolved Brains for each IOI. Note that these data represent number of distinct states in multiple loops, therefore, it is possible for a Brain to visit more states than the IOI. Note also that in traversing the loop once (in one period of the sequence) it is possible to visit some Brain states more than once. For example, the sequence: 6,3,1,1,6,3,1,1,… has a period of 4, but only three distinct states are visited. These results indicate that the number of distinct states visited by evolved Brains, i.e., the size of the state space used to encode temporal information, starts to plateau for longer IOIs.

**Supplementary Figure 3.**
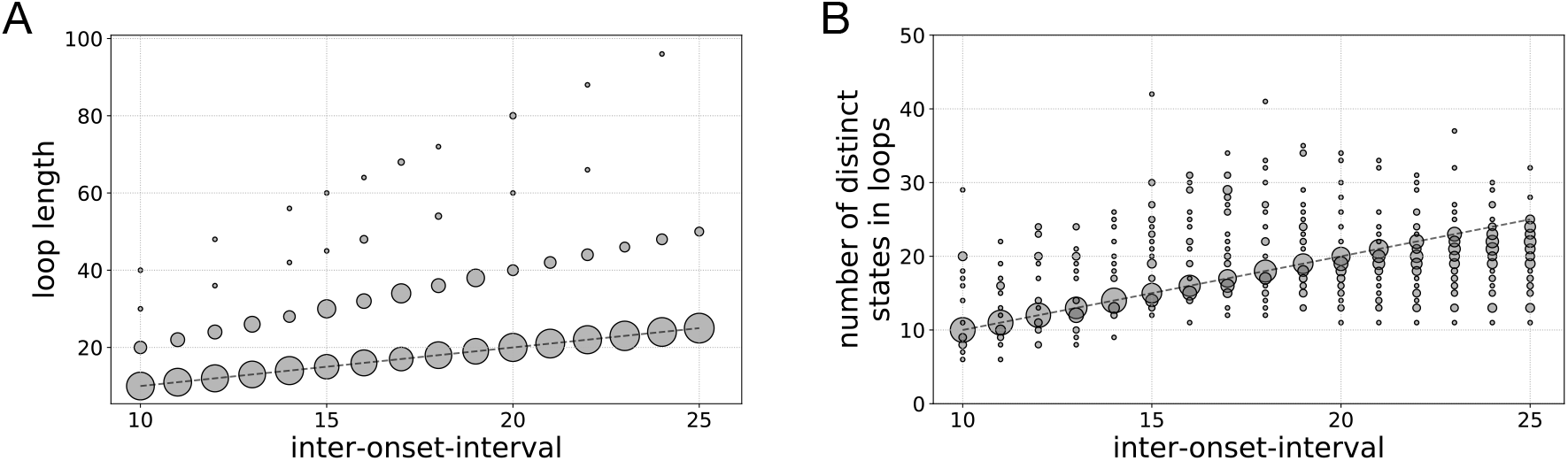
(A) The distribution of loop sizes of 50 evolved brain for each inter-onset-interval (IOI). The size of the markers is proportional to the number of Brains (out of 50) that evolve a particular loop lengths in each IOI. (B) The distribution of number of distinct states in loops visited by Markov Brains in a sequence of rhythmic standard tones, as a function of IOI. The dashed line shows the identity function line.

The duration judgement task in trials with longer IOIs and standard tones is inherently more difficult (see Supplementary Fig. 1B) for two reasons. First, longer rhythms and durations require more memory and computations to encode temporal information, and second, the number of possible oddball tones (in range [1, *IOI* − 1]) is greater in longer IOIs compared to the number of possible oddball tones in shorter IOIs. As a result, Markov Brains need to use progressively larger regions of their state-space to encode the temporal information and moreover, they need more evoltionary time to learn a larger number of patterns; however, state-space size does not grow linearly with IOI but rather begins to plateau (Supplementary Fig. 3B) which, in turn, leads to less accurate performance in duration judgements in trials with longer rhythms and a systematic increase in PSE and CE values. This plateau in utilisation of state-space occurs not because of limitations in Markov Brains capacity but due to historical contingencies in the evolution. More specifically, the fitness landscape is structured in such a way that Markov Brains evolve to perform the duration judgement task for shorter IOIs earlier during the evolutionary course. As a consequence, algorithms that emerge later in evolution that perform the task in longer IOIs are built upon those algorithms evolved earlier. In order to provide further support for the claims we made here, we conducted a series of additional experiments. In the following sections we present results for the evolution of Markov Brains performing duration judgement for various experimental setups that differ slightly from the original experimental setup used in the main text.

#### Longer evolutionary time does not resolve systematic behavioural distortions in longer rhythms/standard tones

In the first set of additional experiment, we continued running the experiments presented in main text (which were run for 2,000 generations) for longer evolutionary time, namely 10,000 generations. Supplementary Fig. 4 shows the fitness values of the best performing agents averaged across 50 runs as a function of IOI and colour-coded at different evolutionary times. We observe that the average fitness values increase in all IOI and standard tones with evolution, however, we still observe the same pattern that the performance drops as IOI increases. Supplementary Supplementary Fig. 5 shows CE values as a function of (IOI, standard tone) at different evolutionary time points. These results show that constant errors in longer IOIs decrease with evolutionary time, however, this decrease slows down considerably and more importantly, a similar trend in CE values vs. (IOI, standard tone) is observed in all generations.

**Supplementary Figure 4.**
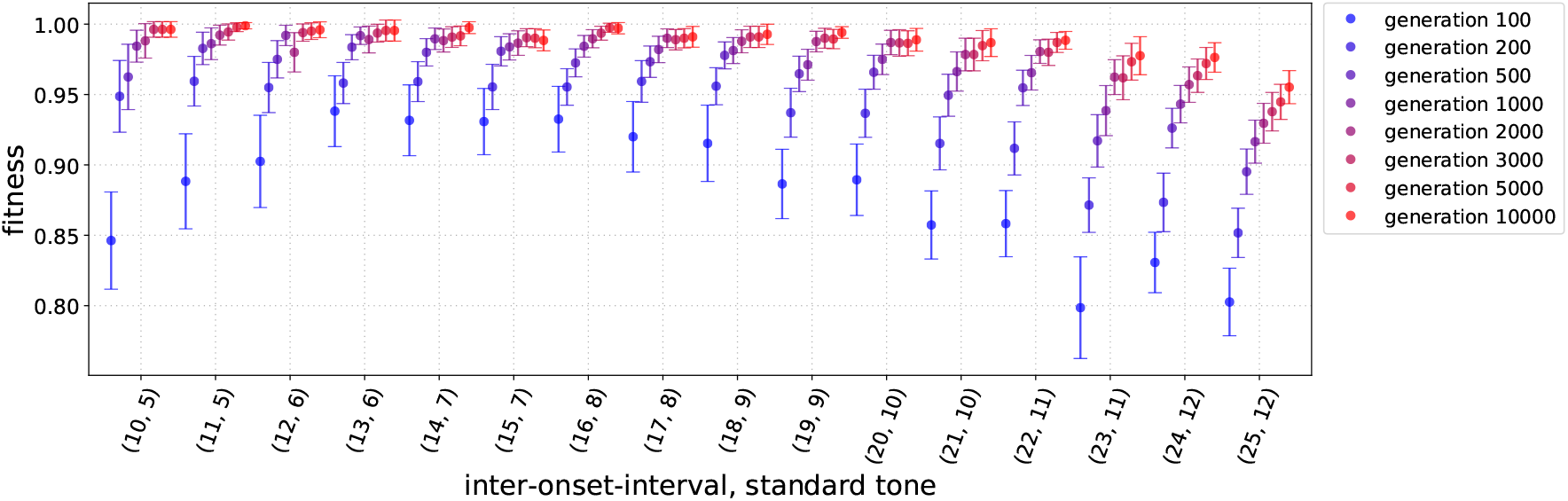
(A) Mean fitness across all 50 lineages and 95% confidence interval color-coded at different evolutionary times as a function of inter-onset-interval, standard tone.

Supplementary Fig. 6 shows the number of distinct states used to encode temporal information corresponding to each IOI at different evolutionary time points. After 100 generations, the distributions of state-space size in shorter rhythms (IOIs 10-14) peak at the IOI (the identity function shown with dashed line) but as the IOI increases the peak of the distribution start to deviate from the identity line and begin to spread more widely. As evolution progresses, the distribution of distinct states in a larger number of IOIs peaks at the identity function but in all the plots shown in Supplementary Fig. 6 (after different number of generations), the distributions that deviate from the identity line correspond to the longest IOIs. For example, after 2,000 generations the distributions for IOIs 23-25 are further from the identity line, and after 10,000 elapsed generations this occurs for IOIs 24, and 25. Recall that we observed a similar pattern in CE values, where at the beginning of evolution CEs for shorter IOIs are around 0 but begin to deviate from 0 for longer IOIs, and as populations evolve further CEs for larger and larger number of IOIs approach 0. Note that the size of the state-space corresponding to each rhythm is indicative of how accurately the representation of that rhythm is encoded in the Brain. And clearly, in longer IOIs Markov Brains do not use as accurate an encoding and therefore, their performance drops for longer IOIs and CE values start to increase systematically.

Here we investigate in more depth the correlation between CE values and the size of state-space used by Markov Brains to encode temporal information. As discussed before, the optimum number of distinct states used to encode stimuli characteristics is the length of the rhythm, i.e., IOI. When the number of distinct states used to encode the rhythm length is smaller than IOI, it means that different time points during that interval have the same representation in the Brain because the Brain must visit some state(s) more than once (at different time points). For example, consider a Brain that is entrained to a rhythm and is traversing a loop in state-space. An oddball tone results in the Brain exiting that loop (we showed such an example in the main text). In this case, if the exit from the loop occurs from a repeated state in that loop, the Brain’s experiences of oddballs that end at different time points would be exactly the same. Alternatively, when the number of distinct states visited when traversing the loops is greater than IOI, it means that the period of that loop is not IOI but a multiple of the IOI. This may also result in less accurate performance in duration judgement task, for example in the judgement of oddballs with the same duration that occur in different positions (recall that oddball tones can occur at 5^*th*^, 6^*th*^, 7^*th*^, or 8^*th*^ position).

In Supplementary Fig. 3B, we observed that the distribution of number of distinct states in loops peaks at IOI for shorter IOIs at the outset of evolution and increasingly more distributions move towards the IOI and accumulate around IOI. Let 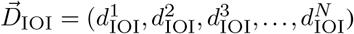, where 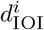 represents the number of distinct states the *i^th^* Brain uses in its loops for a particular IOI, and *N* = 50 since we have 50 evolved Brains. Thus, each distribution in Fig 6 can be represented by a vector 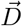. We now calculate the distance of each distribution to the IOI by:

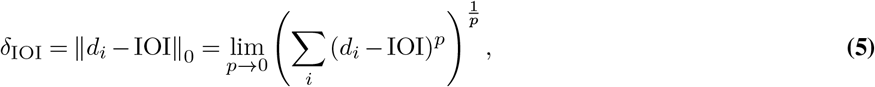

in which ǁǁ_0_ denotes the 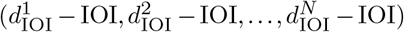. In fact, *δ*_IOI_ simply reflects how many of the 50 Brains do not use exactly IOI distinct states in their loops. We calculated *δ*_IOI_ for each IOI and at different points in evolutionary time. We then normalised these *δ*_IOI_ by the maximum *δ*_IOI_ value. Fig 7 shows absolute CE values as a function of normalised *δ*_IOI_. Each data point shown in grey represents *δ*_IOI_ calculated in a distribution at a specific evolutionary time and a particular IOI in Supplementary Fig. 6).

**Supplementary Figure 5.**
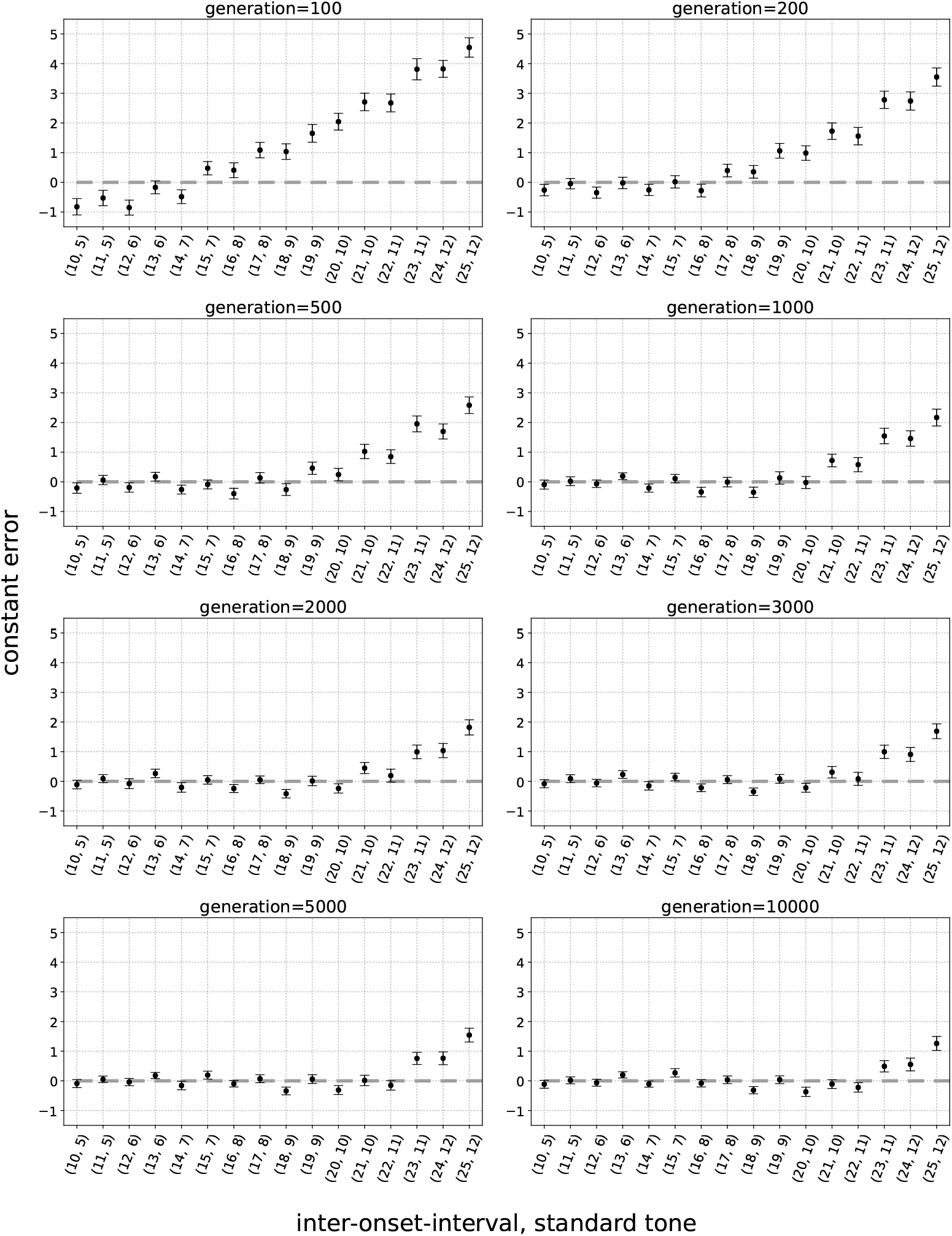
Constant errors and their 95% confidence interval for 50 best performing Brains as a function of inter-onset-interval, standard tone at different evolutionary times. Dashed line shows zero constant error.

**Supplementary Figure 6.**
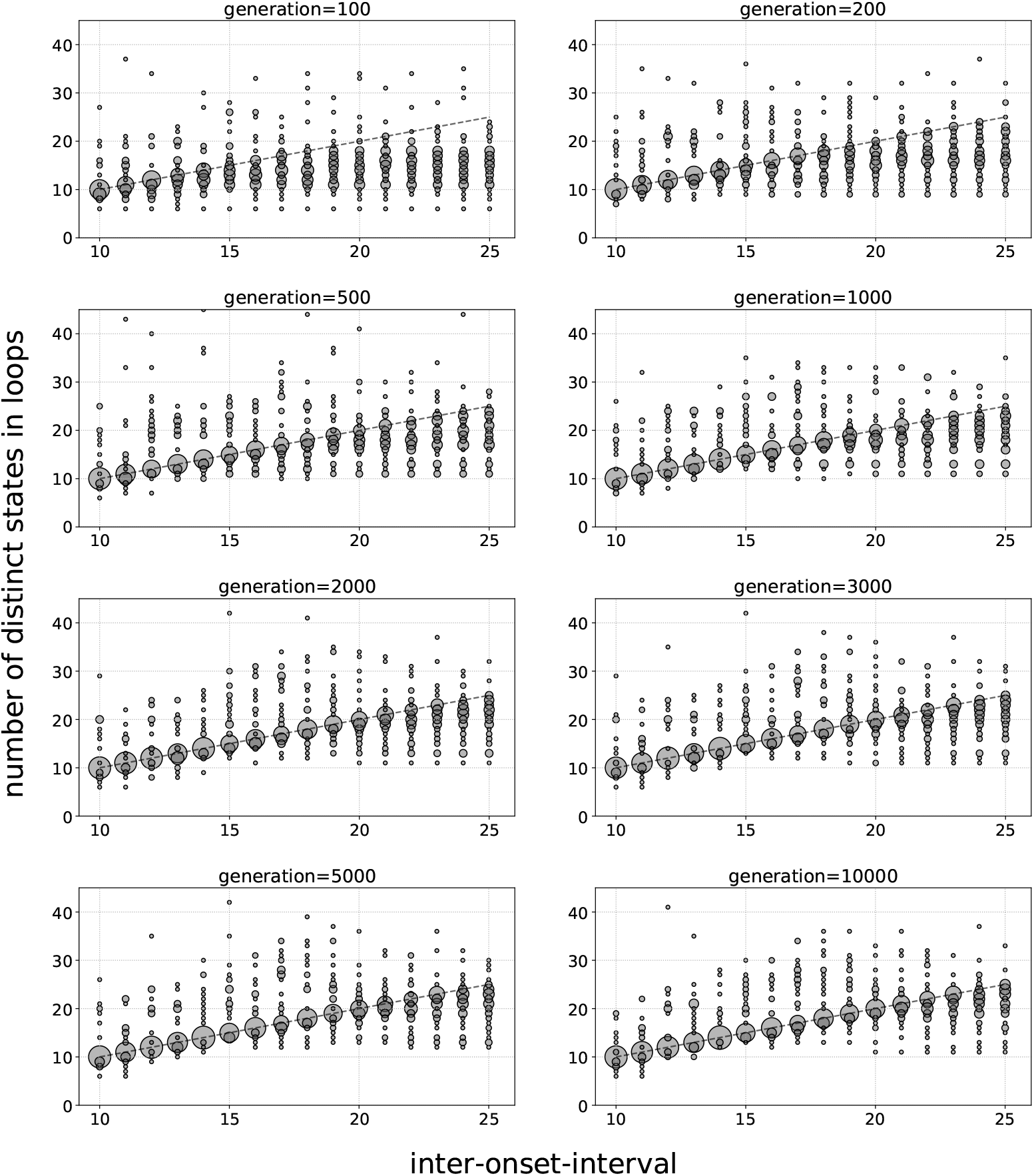
The distribution of number of distinct states used to encode rhythm and standard tone duration, i.e., the number of distinct states in each loop, as a function of inter-onset-interval at different evolutionary times. The size of the circle is proportional to the likelihood at that loop size. The dashed line shows the identity function.

**Supplementary Figure 7.**
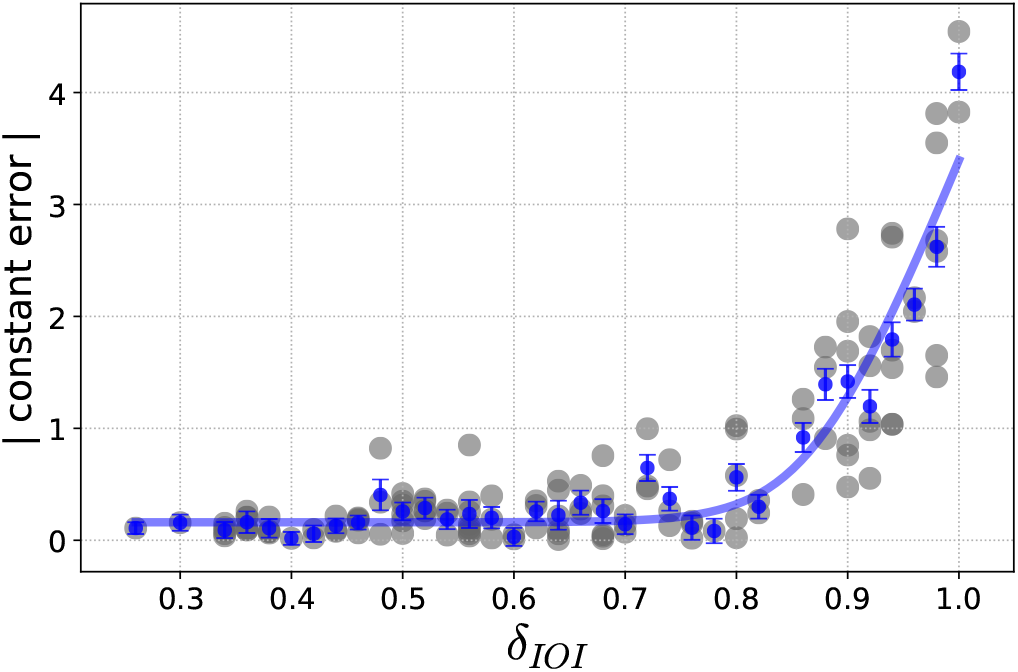
Absolute constant errors (CE) shown in grey as a function of *δ*_IOI_, as well as the binned data and the fitted softplus curve.

**Supplementary Table 1.**
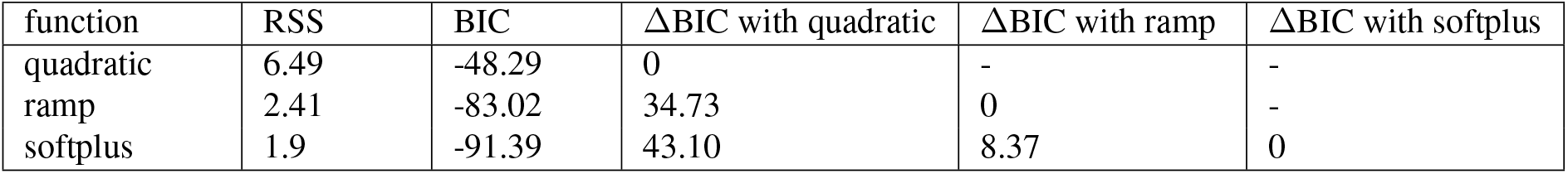
Non-linear regression analysis used to explain the correlation between the constant errors (CE) and *δ*_IOI_ which is a function of the distinct number of states used in encoding stimuli. Residuals sum of squares (RSS), and the Bayesian information criterion. A BIC difference *>* 10 provide very strong support for one model over the other (4).

| function | RSS | BIC | ΔBIC with quadratic | ΔBIC with ramp | ΔBIC with softplus |
| --- | --- | --- | --- | --- | --- |
| quadratic | 6.49 | -48.29 | 0 | - | - |
| ramp | 2.41 | -83.02 | 34.73 | 0 | - |
| softplus | 1.9 | -91.39 | 43.10 | 8.37 | 0 |

We used a non-linear regression analysis (3) to find the correlation between the CE and *δ*_IOI_. Since a large number of data points fall around CE=0 and in the lower range of *δ*_IOI_ (which is not surprising since most trials result in CEs that are not significantly different from 0), we applied binning with constant bin size to this data. Mean values of binned data and their standard deviations as well as the fitted function are also shown in Supplementary Fig. 7. We tested three different kernel functions for regression analysis: 1) quadratic function, 2) ramp function, 3) softplus function (*f* (*x*) = *log*(1 + *e^x^*), which is a differentiable approximation of ramp function). Supplementary Table 1 shows the regression analysis results for three different kernel functions. We compare these three models using Bayesian information criterion (BIC) (4). These results show that the softplus function describes the pattern in the data better than quadratic and ramp function. This pattern can be interpreted as: there is no significant change in CE values for a range of small *δ*_IOI_s, however, by further increasing *δ*_IOI_, at some threshold CEs start to increase linearly with *δ*_IOI_.

#### Training Markov Brains equally in all IOIs and standard tones has a minor effect on behavioural deviations in longer rhythms

In this experimental setup, we used the same set of inter-onset-intervals, standard tones, and oddball tones as used in original experimental setup. The only difference is that the number of evaluations for each (IOI, standard tone) is not constant anymore (in the original setup we evaluate Brains in 22 trials for each IOI, standard tone) but in this modified setup it increases with IOI linearly. Supplementary Table 2 shows the number of evaluation trials as well as IOI, standard tone, and total number of trials for each (IOI, standard tone). Note that we tried to keep the total number of evaluations in this setup, 368 (37.1% of all possible trials), as close as possible to that of the original setup 352 (35.5% of all possible trials). Note also that the number of evaluations in each (IOI, standard tone) is chosen proportionate to the number of oddball tones in that (IOI, standard tone).

Supplementary Fig. 8 shows CE values for this experimental setup as a function of (IOI, standard tone) at different evolutionary time points in the experiments. It is evident that the same trend in CE values that was observed in the original setup can be seen in these experiments too. In particular, after 2,000 generations CEs for (IOI, standard tone)={(23, 11), (24, 12), (25, 12)} are significantly different from 0 and similarly, after 10,000 generations the CE for (25, 12) is significantly different from 0. Supplementary Fig. 9 shows state-space sizes as a function of IOI at different evolutionary time points. Similar to trends observed in the original setup, state-space sizes plateau as IOIs increase and again, their distributions are slightly closer to the identity function (dashed line) but not significantly so. Thus, we conclude that having the same training set size for all IOIs has little to do with distorted behaviours in longer rhythms. Supplementary Fig. 10 shows the binned CE values as a function of *δ*_IOI_ as well as the fitted softplus function. We performed the non-linear regression analysis described before for this experiment and the results are presented in Supplementary Table 3. Similar to previous experiment, the softplus function describes the pattern in CE values and *δ*_IOI_ better than the other two models.

**Supplementary Figure 8.**
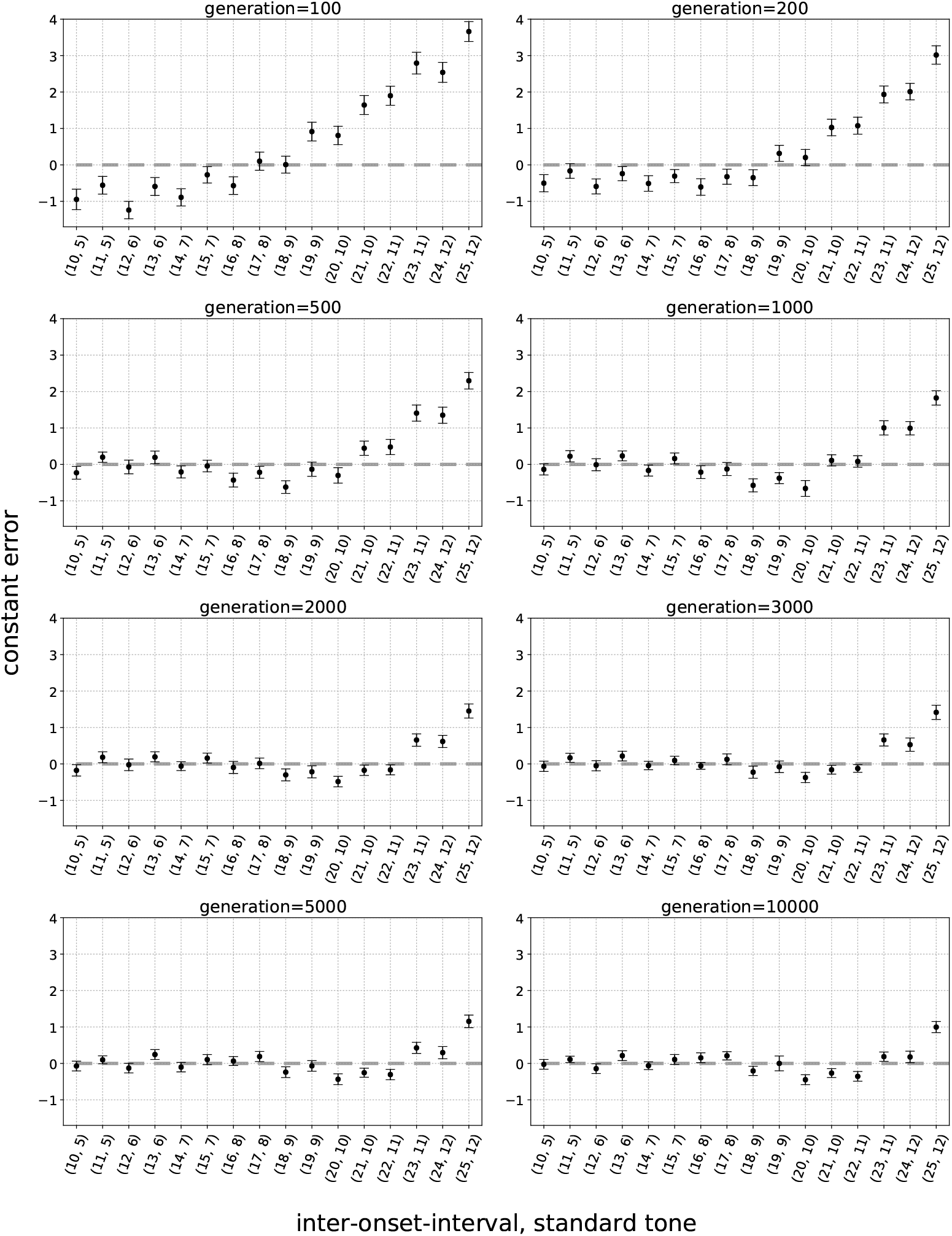
Constant errors and their 95% confidence interval for 50 best performing Brains as a function of inter-onset-interval, standard tone at different evolutionary times. Dashed line shows zero constant error.

**Supplementary Figure 9.**
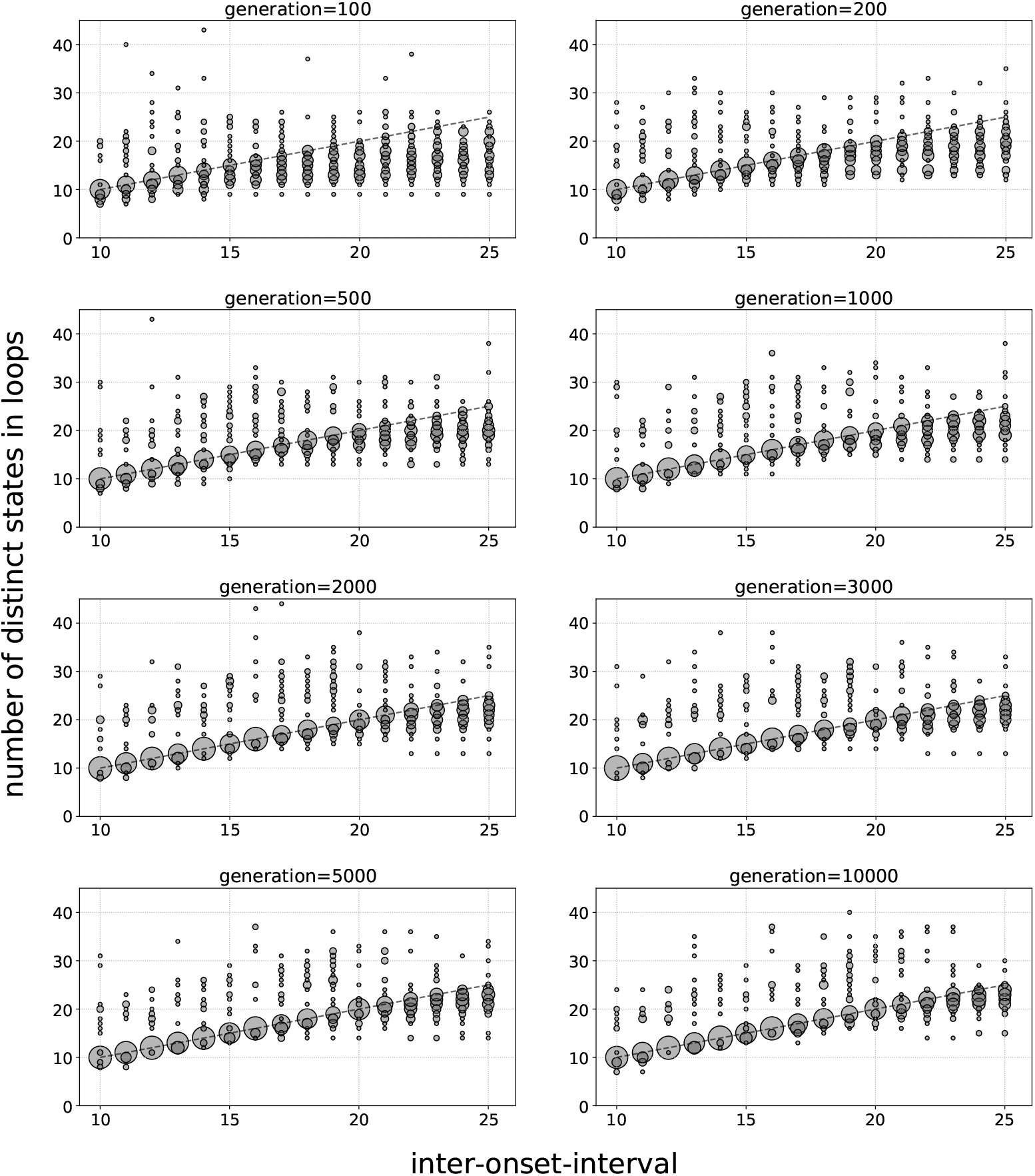
The distribution of number of distinct states used to encode rhythm and standard tone duration, i.e., the number of distinct states in each loop, as a function of inter-onset-interval at different evolutionary times. The dashed line shows the identity function.

**Supplementary Figure 10.**
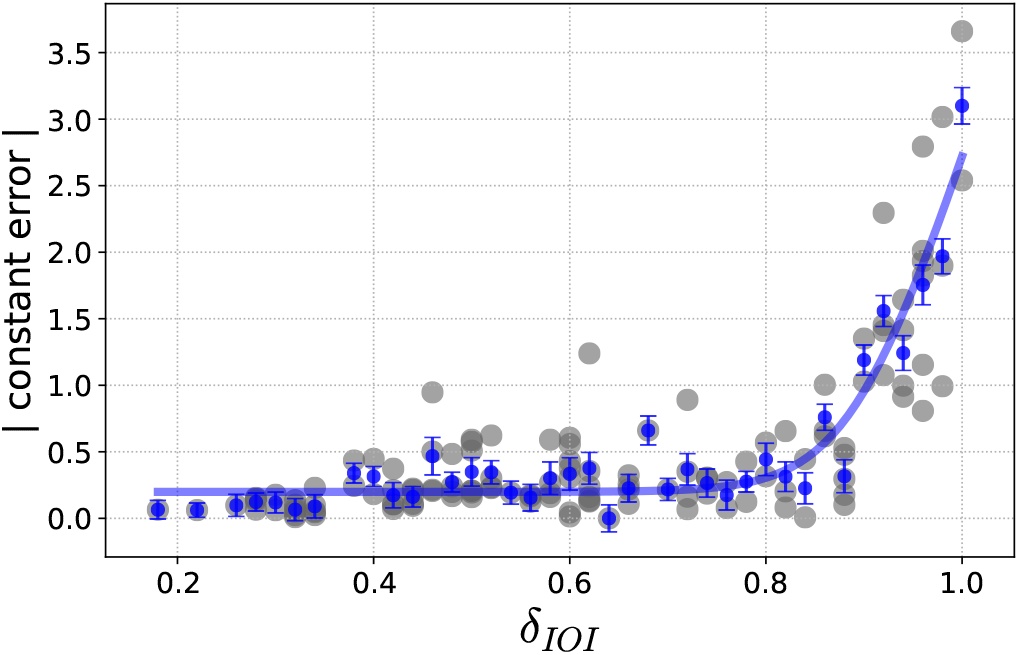
Absolute constant errors (CE) shown in grey as a function of *δ*_IOI_ as well as the binned data and the fitted softplus curve.

**Supplementary Table 2.** Complete set of all inter-onset-intervals, standard tones, and oddball durations used for evolution of duration judgement task. Oddballs can occur in either of 5th, 6th, 7th, or 8th position in the rhythmic sequence. Also, oddball durations are always either shorter or longer than the standard tone.

| Inter-onset-interval | Standard tone duration | Oddball tone duration | Total number of possible trials | number of evaluation trials |
| --- | --- | --- | --- | --- |
| 10 | 5 | {1, 2, 3, 4} , {6, 7, 8, 9} | 32 | 8 |
| 11 | 5 | {1, 2, ..., 4} , {6, ..., 10} | 36 | 10 |
| 12 | 6 | {1, 2, ..., 5} , {7, ..., 11} | 40 | 12 |
| 13 | 6 | {1, 2, ..., 5} , {7, ..., 12} | 44 | 14 |
| 14 | 7 | {1, 2, ..., 6} , {8, ..., 13} | 48 | 16 |
| 15 | 7 | {1, 2, ..., 6} , {8, ..., 14} | 52 | 18 |
| 16 | 8 | {1, 2, ..., 7} , {9, ..., 15} | 56 | 20 |
| 17 | 8 | {1, 2, ..., 7} , {9, ..., 16} | 60 | 22 |
| 18 | 9 | {1, 2, ..., 8} , {10, ..., 17} | 64 | 24 |
| 19 | 9 | {1, 2, ..., 8} , {10, ..., 18} | 68 | 26 |
| 20 | 10 | {1, 2, ..., 9} , {11, ..., 19} | 72 | 28 |
| 21 | 10 | {1, 2, ..., 9} , {11, ..., 20} | 76 | 30 |
| 22 | 11 | {1, 2, ..., 10} , {12, ..., 21} | 80 | 32 |
| 23 | 11 | {1, 2, ..., 10} , {12, ..., 22} | 84 | 34 |
| 24 | 12 | {1, 2, ..., 11} , {13, ..., 23} | 88 | 36 |
| 25 | 12 | {1, 2, ..., 11} , {13, ..., 24} | 92 | 38 |

**Supplementary Table 3.** Non-linear regression analysis used to explain the correlation between the constant errors (CE) and *δ*_IOI_ which is a function of the distinct number of states used in encoding stimuli. Residuals sum of squares (RSS), and the Bayesian information criterion.

| function | RSS | BIC | ΔBIC with quadratic | ΔBIC with ramp | ΔBIC with softplus |
| --- | --- | --- | --- | --- | --- |
| quadratic | 5.36 | -66.42 | 0 | - | - |
| ramp | 1.59 | -113.77 | 47.35 | 0 | - |
| softplus | 1.4 | -118.65 | 52.23 | 4.88 | 0 |

#### Constant errors in longest rhythms are greater than zero regardless of trial size

In order to show that the deviations of PSE (from the point of objective equality, i.e., standard tone) in longer IOI, and standard tones is not specific to a particular value of IOI or standard tone, we used two experimental setups where one has a smaller set of (IOI, standard tone) with shorter IOIs and standard tone durations, and one that has a larger set of (IOI, standard tone) with longer rhythms, standard tones. The first training set is similar to the original experimental setup but we excluded trials with the following inter-onset-intervals and standard tones from the original setup: {(23,11), (24,12), (25,12)}. Similar to the original setup, oddball tones can vary from 1 to IOI-1. In this experimental setup, there are 728 possible trials and all agents are evaluated in 20 trials from each IOI and standard tone (10 with longer and 10 with shorter oddball tones) which is 35.7% of all possible trials (in the original setup evaluation trials set was 35.5% of all possible trials).

Supplementary Fig. 11 shows mean constant errors as a function of standard tones at different evolutionary times for this experimental setup. The increase in CEs is again observed for longer IOIs and noticeably, after 2000 generations in trials with (IOI, standard tone)={(10, 5), (11,5)}, all 50 Brains perform the duration judgement task perfectly (100% performance for all oddball tones in those rhythms) and we observe Brains perform the duration judgement task perfectly in more IOIs, and standard tone in later generations, for example after 10,000 generations Brains perform perfectly in (IOI, standard tone)={(10, 5), (11, 5), (12, 6), (14,7)}. Cognitive scientists and psychophysicists are not in general interested in “trivial” experiments in which all the subjects answer 100% of questions correctly; therefore, we did not design our experimental setup such that Brains evolve to achieve 100% fitness either. Supplementary Fig. 12 shows state-space size distributions as a function of IOI for different evolutionary time points. It is again evident that the state-space sizes start to plateau for longer IOIs but of course, not as drastically as in the original setup. The CE values, as well as binned means and their standard deviations, are shown as a function of *δ*_IOI_ are shown in Supplementary Fig. 13. In Supplementary Fig. 13, the blue dashed line shows the fitted softplus function. The results of the non-linear regression analysis are shown in Supplementary Table 4. We again observe that the softplus function describes the pattern in CE values and *δ*_IOI_ better than the other two functions.

The second experimental setup has all the trials from the original and we also added the following inter-onset-intervals and standard tones: {(26,13), (27,13), (28,14), (29,14)}. In this experimental setup, there are 1400 possible trials and all agents are evaluated in 24 trials from each IOI and standard tone (12 with longer and 12 with shorter oddball tones) which is 34.3% of all possible trials to maintain the same ratio of evaluation trials to all possible trials. Supplementary Fig. 14 shows mean constant errors as a function of standard tones at different evolutionary times for this experimental setup. These results show a similar pattern in CE values and more importantly, we observe that the CEs for the inter-onset-interval and standard tones {(23,11), (24,12), (25,12)} are not significantly different from 0 whereas in the original experiment, CEs were significantly different from 0 in the same trials, i.e., {(23,11), (24,12), (25,12)}. Supplementary Fig. 15 shows state-space size distributions as a function of inter-onset-intervals for different evolutionary time points. We again observe that the state-space sizes start to plateau for longer IOIs but of course, but not as drastically as in the original setup. We performed the non-linear regression analysis on these data as well and the results are shown in Supplementary Table 5. As observed in previous results, the softplus function describes the pattern in CE values and *δ*_IOI_ better than the other two models. The CE values, the binned data mean and standard deviations, and the fitted softplus function is shown in Supplementary Fig. 16.

**Supplementary Figure 11.**
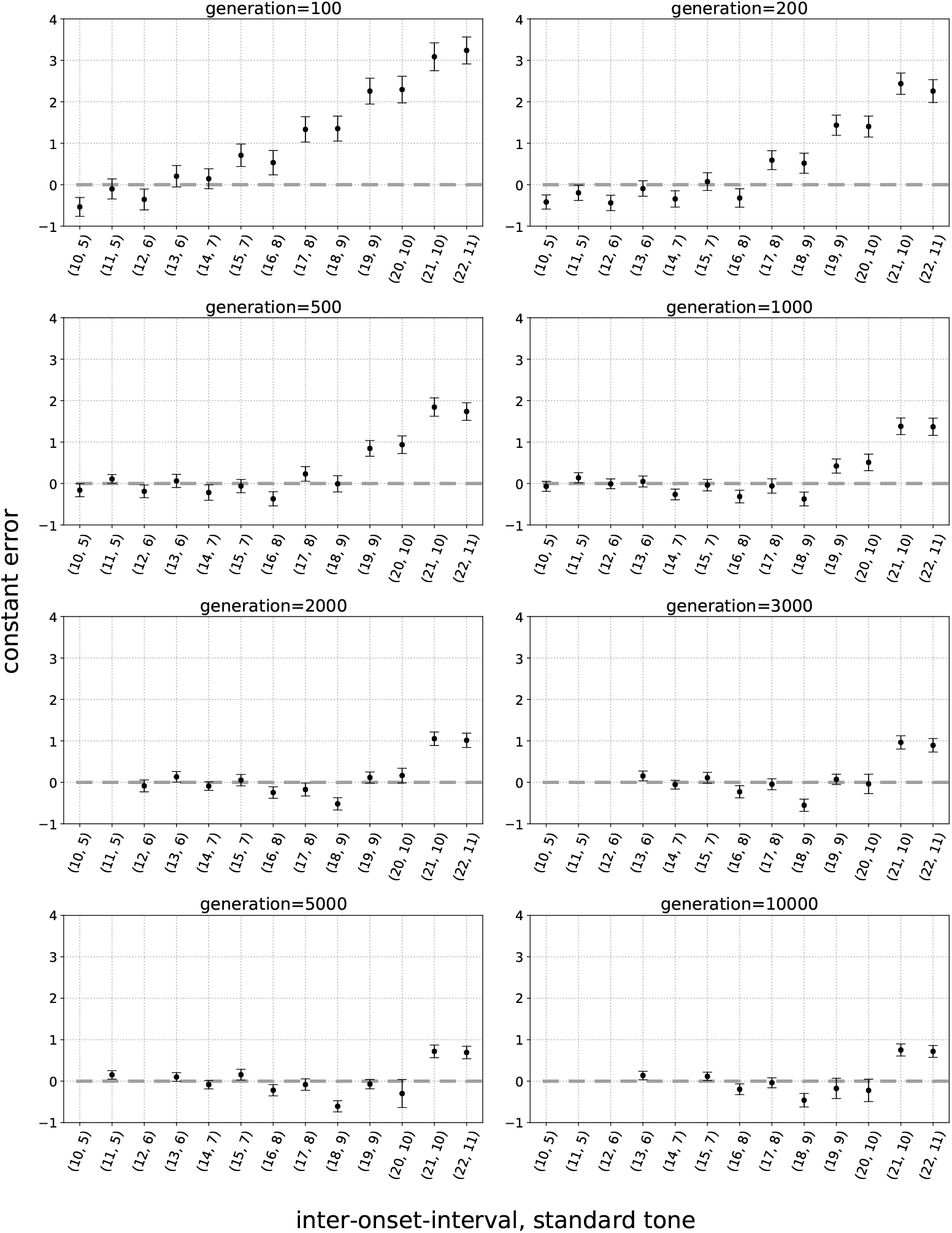
Constant errors and their 95% confidence interval for 50 best performing Brains as a function of inter-onset-interval, standard tone at different evolutionary times. There are some missing data points in these plots which is due to the fact that in those trials the performances of all 50 Brains are 100%, as a result, PSE would be exactly equal to the standard tone and the slope of the psychometric function would be infinity. Dashed line shows zero constant error.

These results reaffirm that the entrainment and duration judgement task become much more difficult for longer (IOIs, standard tone) and with greater set of trials, and that furthermore, Markov Brains do have the capacity to use greater regions of the state-space and perform more accurately in longer IOIs. However, the historical contingencies in such fitness landscapes lead to less accurate strategies in duration judgements in longer IOIs which results from using smaller regions in state-space.

**Supplementary Figure 12.**
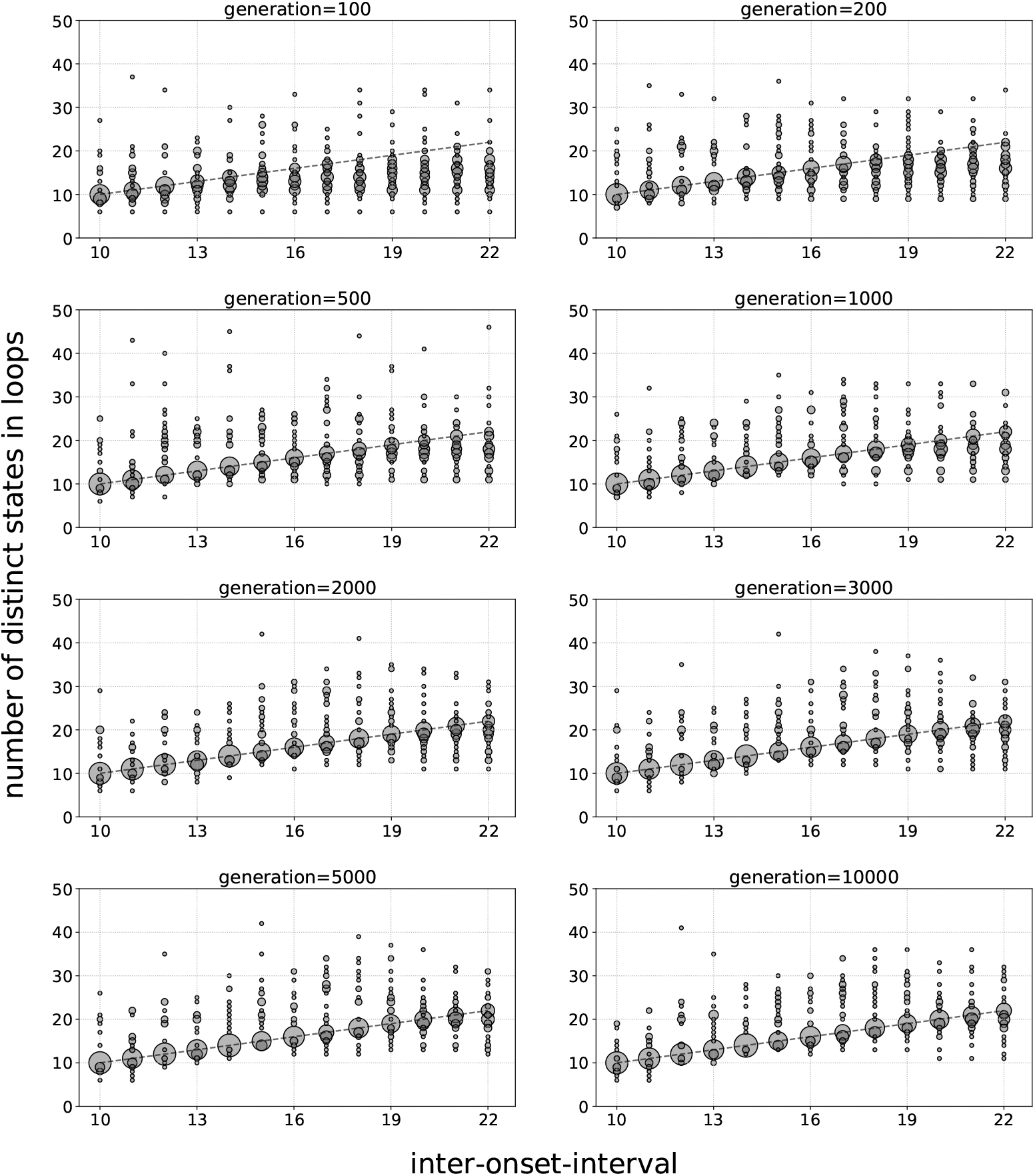
The distribution of number of distinct states used to encode rhythm and standard tone duration, i.e., the number of distinct states in each loop, as a function of inter-onset-interval at different evolutionary times. The dashed line shows the identity function.

**Supplementary Figure 13.**
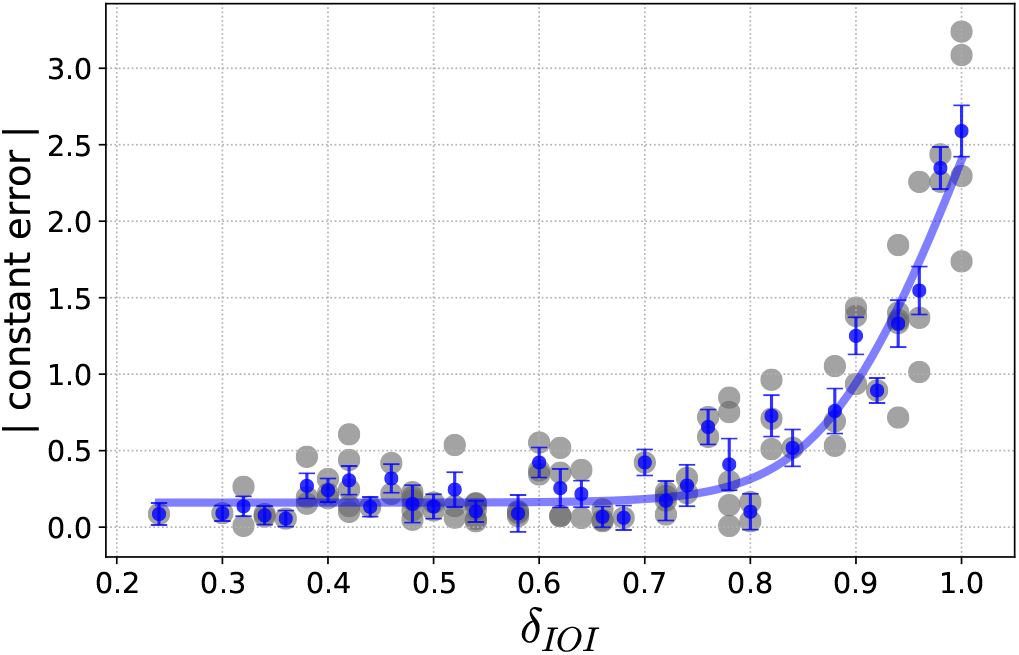
Absolute constant errors (CE) shown in grey as a function of *δ*_IOI_, as well as the binned data and the fitted softplus curve.

**Supplementary Table 4.** Non-linear regression analysis used to explain the correlation between the constant errors (CE) and *δ*_IOI_ which is a function of the distinct number of states used in encoding stimuli. Residuals sum of squares (RSS), and the Bayesian information criterion.

| function | RSS | BIC | $\Delta\text{BIC}$ with quadratic | $\Delta\text{BIC}$ with ramp | $\Delta\text{BIC}$ with softplus |
| --- | --- | --- | --- | --- | --- |
| quadratic | 2.91 | -76.33 | 0 | - | - |
| ramp | 1.48 | -99.97 | 23.64 | 0 | - |
| softplus | 0.98 | -114.51 | 38.18 | 14.54 | 0 |

**Supplementary Figure 14.**
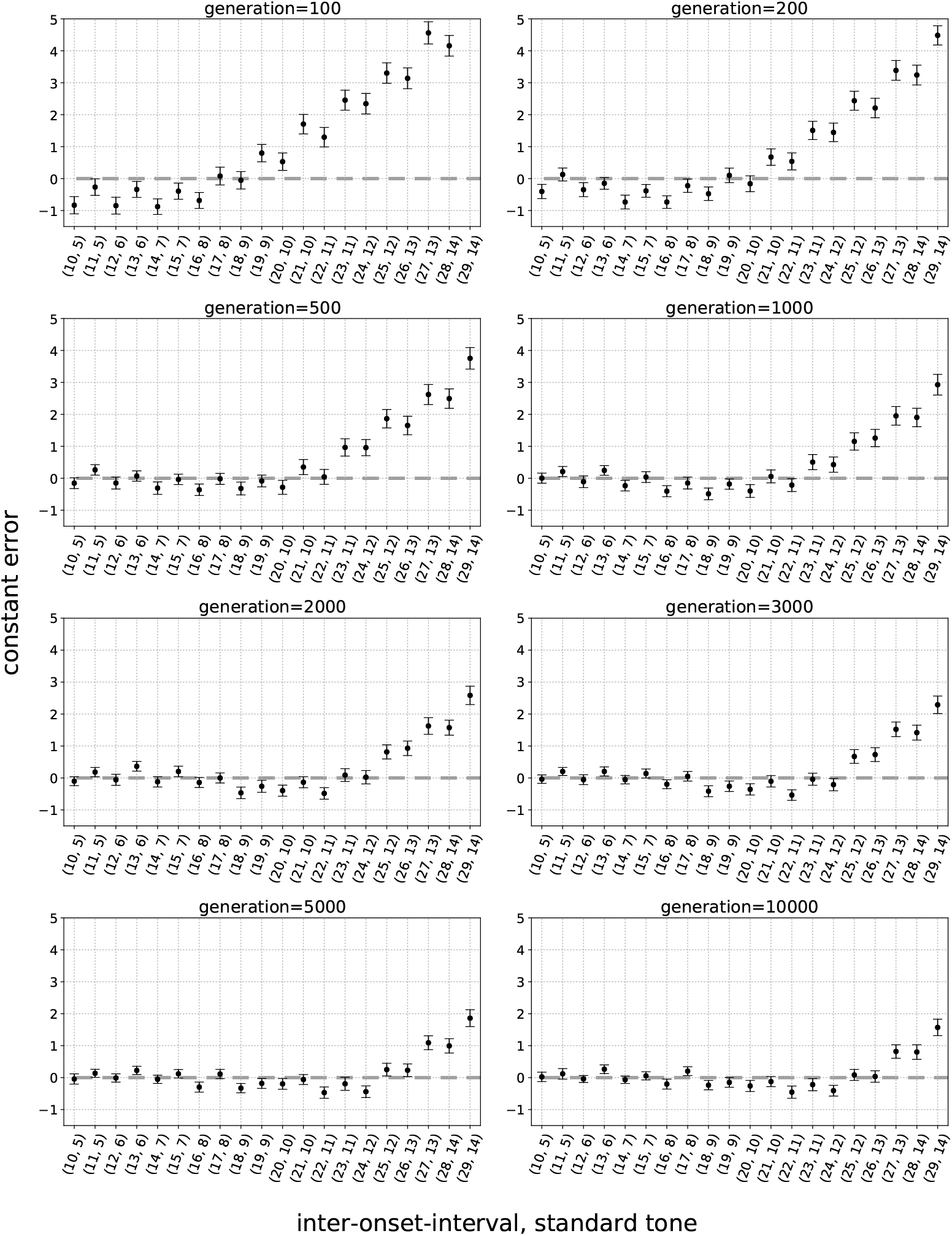
Constant errors and their 95% confidence interval for 50 best performing Brains as a function of inter-onset-interval, standard tone at different evolutionary times. There are some missing data points in these plots which is due to the fact that in those trials the performances of all 50 Brains are 100%, as a result, PSE would be exactly equal to the standard tone and the slope of the psychometric function would be infinity. Dashed line shows zero constant error.

**Supplementary Figure 15.**
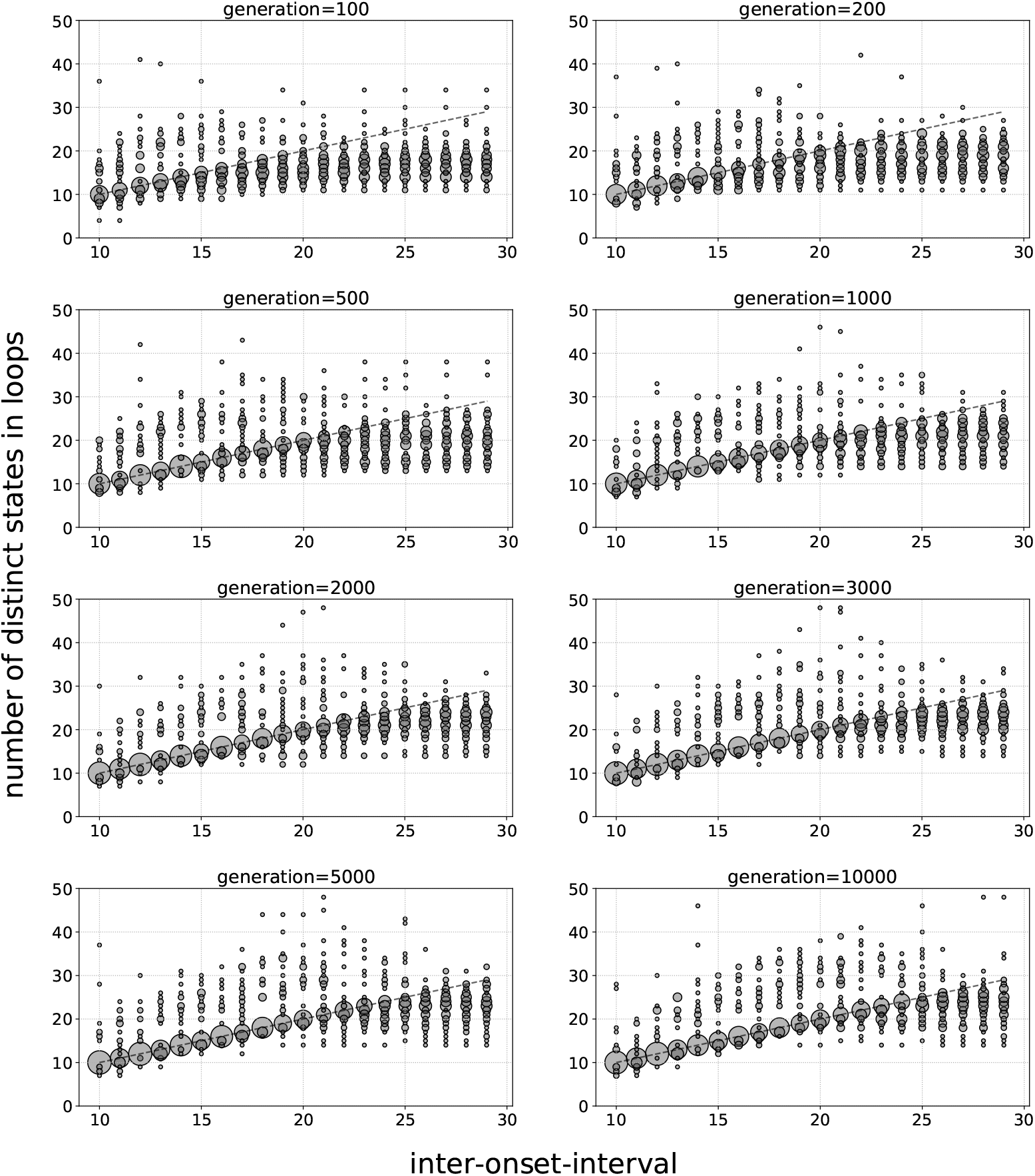
The distribution of number of distinct states used to encode rhythm and standard tone duration, i.e., the number of distinct states in each loop, as a function of inter-onset-interval at different evolutionary times. The dashed line shows the identity function.

**Supplementary Figure 16.**
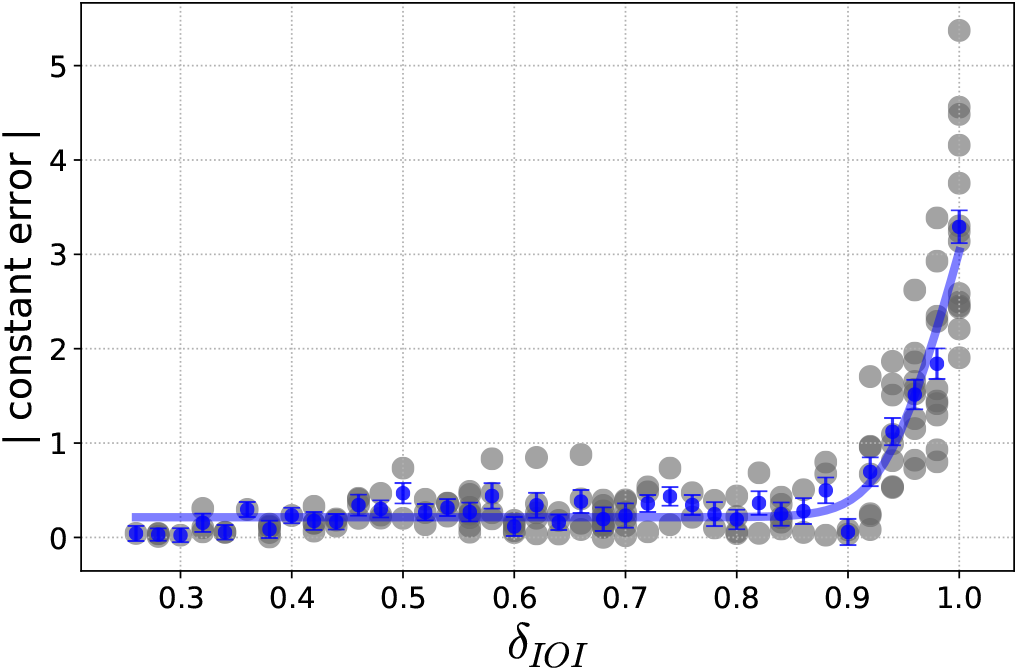
Absolute constant errors (CE) shown in grey as a function of *δ*_IOI_, as well as the binned data and the fitted softplus curve.

**Supplementary Table 5.** Non-linear regression analysis used to explain the correlation between the constant errors (CE) and *δ*_IOI_ which is a function of the distinct number of states used in encoding stimuli. Residuals sum of squares (RSS), and the Bayesian information criterion.

| function | RSS | BIC | ΔBIC with quadratic | ΔBIC with ramp | ΔBIC with softplus |
| --- | --- | --- | --- | --- | --- |
| quadratic | 7.03 | -53.20 | 0 | - | - |
| ramp | 1.03 | -126.34 | 73.14 | 0 | - |
| softplus | 0.89 | -131.92 | 78.72 | 5.58 | 0 |

**Supplementary Video S1.** Animation showing state transitions of Markov Brain shown in figure S2 entraining to the rhythm with IOI=10, and std tone=5.

**Supplementary Video S2.** Animation showing state transitions of Markov Brain shown in figure S2 judging a longer oddball tone IOI=10, std tone=5, and oddball tone =6.

**Supplementary Video S3.** Animation showing state transitions of Markov Brain shown in figure S2 judging a shorter oddball tone IOI=10, std tone=5, and oddball tone =4.

**Supplementary Video S4.** Animation showing state transitions of Markov Brain shown in figure S2 judging a shorter oddball tone that is late 2 time steps, IOI=10, std tone=5, oddball tone =4, and onset=+2.

**Supplementary Video S5. Animation showing state transitions of Markov Brain shown in figure S2 judging a shorter oddball tone that is early 2 time steps, IOI=10, std tone=5, oddball tone =6, and onset=-2.**

## Footnotes

1 Here and below, to avoid confusion we use “Brain” with a capital B to denote artificial brains, while biological brains remain just “brains”.

## Notes

### Competing Interest Statement

The authors have declared no competing interest.

### Summary of Updates

Title changed. Small edits.

